# Resistance of *Dickeya solani* strain IPO 2222 to lytic bacteriophage vB_Dsol_D5 (ΦD5) results in fitness tradeoffs for the bacterium during infection

**DOI:** 10.1101/2022.02.23.481671

**Authors:** Przemyslaw Bartnik, Kinga Lewtak, Marta Fiołka, Paulina Czaplewska, Magdalena Narajczyk, Robert Czajkowski

## Abstract

Resistance to bacteriophage infections protects bacteria in phage-full environments, allowing them to survive and multiply in the presence of their viral predators. However, such resistance may cause direct costs for strains linked with the ecological fitness expressed as reduced competitiveness for resources or reduced virulence or both. Unfortunately, limited knowledge exists about such costs paid by phage-resistant plant pathogenic bacteria in their natural environments. This study analyzed the costs of phage resistance paid by broad host phytopathogenic pectinolytic bacterium *Dickeya solani* both *in vitro* and in potato (*Solanum tuberosum* L.) plants. Thirteen *D. solani* IPO 2222 Tn5 mutants were identified that exhibited resistance to infection caused by lytic bacteriophage vB_Dsol_D5 (ΦD5). The genes disrupted in these 13 mutants encoded proteins involved in the synthesis of the bacterial envelope components (viz. LPS, EPS and capsule). The ability of ΦD5-resistant *D. solani* mutants to colonize and cause symptoms on potato plants as well as other phenotypes that are known to contribute to the ecological fitness of *D. solani* in-plant environment, including growth rate, production of effectors, swimming and swarming motility, use of various carbon and nitrogen sources and biofilm formation were assessed. Although phage resistance did not affect most of the phenotypes of ΦD5-resistant *D. solani* evaluated *in vitro*, all phage resistant mutants were significantly compromised in their ability to survive on and colonize and cause disease symptoms in potato plants. This study is, to our knowledge, one of few to show the direct link between phage resistance and the fitness of plant pathogenic bacteria and the first one to assess phage-host associations for *D. solani*.

## Introduction

Bacteriophages (bacterial viruses) are the most abundant biological entities in the biosphere. With their total estimated number of ca. 10^31^ phage particles, they are the main driving force of bacterial adaptation and evolution (Cheetham and Katz, 1995; Campbell, 2003; Mann, 2005) (Thurber, 2009). Likewise, bacterial viruses play a critical role in maintaining bacterial diversity in the environment (Buckling and Rainey, 2002; Forde et al., 2008; Dennehy, 2012). It is estimated that even up to 20% of bacterial cells are killed daily due to phage infections worldwide (Suttle, 1994). However, surprisingly, this enormous mortality of bacterial cells does not lead to the global disappearance of sensitive host populations or the development of resistant ones taking over the given ecological niche (Chibani-Chennoufi et al., 2004; Duffy and Forde, 2009; Koskella and Brockhurst, 2014). On the contrary, phage-susceptible and phage-resistant bacterial populations are frequently reported to coexist both in natural (e.g. ocean, phyllosphere)(Waterbury and Valois, 1993; Koskella and Parr, 2015) and human-controlled and engineered environments (e.g. wastewater treatment facilities, agricultural fields)(Hantula et al., 1991; Jones et al., 2007; Fernandez et al., 2018).

It is hypothesized that the stable coexistence of phage-resistant and phage-susceptible bacterial populations occurs because bacteria pay constant costs of phage resistance despite whether the viruses are present or not in the environment (Stern and Sorek, 2011; Bradde et al., 2017; Burmeister and Turner, 2020; Naureen et al., 2020). Such costs are primarily linked with the altered fitness of resistant bacteria and are usually manifested by reduced competitiveness for resources and/or reduced virulence or both (Labrie et al., 2010; Koderi Valappil et al., 2021). However, it should also be highlighted that not all evolved phage resistance traits suffer costs (Lythgoe and Chao, 2003; Mizoguchi et al., 2003), as well as that the costs may also depend on the environmental context and the mechanism of phage resistance itself (Lennon et al., 2007; Vale et al., 2015).

Even though costs of resistance to phage infection have been reported in different phage-host systems (Abedon, 2012; Keen, 2014; Burmeister and Turner, 2020), relatively little is known about these costs paid specifically by phytopathogenic bacteria residing in their natural habitats, including plant pathogens interactions with lytic bacteriophages in agricultural fields. For example, no broad studies have been done to date to assess the costs of phage resistance in other than *P. parmentieri* Soft Rot *Pectobacteriaceae* (SRP) species.

Soft Rot *Pectobacteriaceae* (SRP bacteria: *Pectobacterium* spp. and *Dickeya* spp.) are an excellent model for studying phage-host interaction and co-adaptation in the environment. These bacteria are considered among the top ten most important agricultural phytopathogens worldwide (Mansfield et al., 2012). SRP cause significant losses in crop production (up to 40%), with disease severity dependent on weather conditions, plant susceptibility and pathogen inoculum (Perombelon, 2002; Charkowski, 2018). *Pectobacterium* spp. and *Dickeya* spp. are widespread in various ecological niches, including natural and agricultural soils, water, sewage, the surface of host and non-host plants, and the surface and interior of insects (Perombelon and Kelman, 1980; Fikowicz-Krosko et al., 2017; Rossmann et al., 2018). Because of the diverse environments in which SRP bacteria can be located, these pathogens apparently also exhibit various lifestyles because of their transfer between these diverse environments: for example, from plant to soil, from plant to plant, from host to non-host plant, from surface/irrigation water to plant and *vice versa* (Charkowski, 2018). In all of these surroundings, the SRP bacteria can encounter lytic bacteriophages and, as a result, may become immediately and repeatedly infected (Batinovic et al., 2019). Under constant high infection pressure, the emergence of phage-resistant SRP variants seems to be inevitable (Wright et al., 2019; Borin et al., 2021). However, it remains unclear what are the ecological fitness costs for SRP bacteria to become resistant to viral infections.

The purpose of this study was to assess the fitness costs paid by phage resistant variants of *D. solani* both *in vitro* and *in planta*. *D. solani* is an emerging plant pathogen causing soft rot disease symptoms in a variety of crops and nonfood crops worldwide (Toth et al., 2011; van der Wolf et al., 2014). The bacterium was first reported in potato in the early 2000s (Czajkowski et al., 2009a) following its establishment as a new *Dickeya* species with strain IPO 2222 as a type strain in 2014 (van der Wolf et al., 2014). Till present, *D. solani* was reported to be present in the majority of European countries (Toth et al., 2011; van der Wolf et al., 2014) as well as in Israel (Tsror Lahkim et al., 2013), Georgia (Tsror et al., 2011), Turkey (Ozturk and Aksoy, 2017) and Brazil (Cardoza et al., 2017) causing increasing losses in crop production. Surprisingly, most of the *D. solani* strains analyzed so far belong to the same halophyte and express only a low number of genetic differences (Khayi et al., 2016).

Our study model involved *D. solani* strain IPO 2222 (van der Wolf et al., 2014) and lytic bacteriophage vB_Dsol_D5 (ΦD5) (Czajkowski et al., 2014a; Czajkowski et al., 2017a). We isolated ΦD5 in 2014 as a broad host lytic bacteriophage able to infect *D. solani* strains as well as strains belonging to three other *Dickeya* species (*D. zeae*, *D. dianthicola* and *D. dadantii*) (Czajkowski et al., 2014a). Phage ΦD5 belongs to genus *Limestonevirus* and family *Ackermannviridae* (Adriaenssens et al., 2012b; Adriaenssens et al., 2012a; Petrzik et al., 2021). From all bacteriophages infecting *D. solani* reported until now, phages belonging to the *Limestonevirus* genus seems to be the most abundant in the environment (Petrzik et al., 2021). Limestoneviruses including ΦD5 have been isolated in various European countries, including Belgium, Poland, Russia and the United Kingdom. They express a high level of genetic homogeneity. Furthermore, bacteriophage vB_Dsol_D5 (ΦD5) has been extensively studied as a biological control agent against *D. solani* in the potato ecosystem (Czajkowski et al., 2017a).

Using Tn5-based random mutagenesis, we aimed to identify and characterize those *D. solani* IPO 2222 transcriptional units (genes and operons) encoding structures required for ΦD5 attachment and infection susceptibility to better understand the molecular determinants responsible for fitness alterations of phage-resistant bacterial variants. Furthermore, we investigated the hypothesis that ΦD5-resistant *D. solani* mutants may be at a fitness drawback to the wild-type IPO 2222 strain and that the level of this tradeoff is dependent on the environmental context in which such resistance occurred.

## Materials and Methods

### Bacteriophages, bacterial strains, and growth media

The lytic bacteriophage vB_Dsol_D5 (□D5) was isolated and characterized in our former studies (Czajkowski et al., 2014a; Czajkowski et al., 2014b; Czajkowski et al., 2017a). For this work, □D5 was propagated on its wild-type host, *D. solani* strain IPO 2222 (van der Wolf et al., 2014), and tittered as described earlier (Czajkowski et al., 2014a). The stock concentration of □D5 phage particles used in all experiments was ca. 10^8^ - 10^9^ plaque-forming units (PFU) mL^-1^ in tryptone soya broth (TSB, Oxoid) or in quarter-strength (1/4) Ringer’s buffer (Merck) unless stated otherwise. The strains used in this study are listed in Supplementary Table 1. The collection of 10,000 *D. solani* Tn5 mutants generated in our former studies (Lisicka et al., 2018; Czajkowski et al., 2020) was used as a source of ΦD5-resistant *D. solani* IPO 2222 mutants. *D. solani* wild type (WT) strain IPO 2222 (van der Wolf et al., 2014) was cultivated for 24-48 h at 28 °C on tryptic soy agar (TSA, Oxoid), in tryptone soy broth (TSB, Oxoid) or M9 minimal medium (MP Biomedicals) supplemented with glucose (Sigma-Aldrich) to a final concentration of 0.4 %. 15 g L^-1^ bacteriological agar (Oxoid) was added to solidify the media. If required, the bacterial media were supplemented with 50 µg mL^-1^neomycin (Sigma-Aldrich), 150 µg mL^-1^ ampicillin (Sigma-Aldrich) or 40 µg mL^-1^tetracycline (Sigma-Aldrich). The liquid bacterial cultures were agitated during incubation (120 rpm). To prevent fungal growth, cycloheximide (Sigma-Aldrich) was added to the growth medium in the final concentration of 200 µg mL^-1^.

### Selection of □D5-resistant D. solani mutants

Phage-resistant *D. solani* Tn5 mutants were preselected as previously described (Czajkowski et al., 2019). To validate resistance to bacteriophage infection, each Tn5 mutant exhibiting a □D5-resistant phenotype in the initial screen was exposed to repeatable phage challenge assays (at least 2 independent assays per mutant) and plaque formation assays (at least two independent assays per mutant) on selected Tn5 mutant host as previously described (Czajkowski et al., 2014a).

### Identification of the transposon insertion sites in phage-resistant D. solani mutants

To precisely localize the Tn5 insertion sites in the genomes of *D. solani* mutants resistant to □D5 infection, the genomes of the selected mutants were sequenced and analyzed. The genomic DNA of each mutant was isolated, sequenced, and assembled into a draft genome at the Laboratory of DNA Sequencing and Oligonucleotide Synthesis (Institute of Biochemistry and Biophysics of the Polish Academy of Science, Warsaw, Poland) using Illumina technology. Structural and functional annotations of the draft Tn5 genomes were acquired from RAST (Rapid Annotation using Subsystem Technology (http://rast.nmpdr.org/) (Aziz et al., 2008). The position of the Tn5 transposons in the draft genomes of *D. solani* IPO 2222 Tn5 mutants was established using BlastN and BlastX alignments accessed *via* the NCBI website (http://blast.ncbi.nlm.nih.gov/Blast.cgi) (Altschul et al., 1990). Using the available complete genome sequence of *D. solani* WT strain IPO 2222 (Genbank accession: CP015137) (Khayi et al., 2016) and the obtained draft Tn5 mutants genomes, the insertion of the Tn5 transposon in the bacterial chromosome was assessed similarly as described earlier (Czajkowski et al., 2020). For each mutant, ca. 1,000-to 5,000-bp-long sequences adjoining the Tn5 transposon site were examined to determine the genomic context of each Tn5 transposon-disrupted gene (Lisicka et al., 2018; Czajkowski et al., 2020). The function of the disrupted genes was inferred using BlastN and BlastX alignments accessed *via* the NCBI website (https://blast.ncbi.nlm.nih.gov/Blast.cgi). Similarly, the functions of any unannotated open reading frames encoding hypothetical proteins or proteins without apparent sequence homology to proteins deposited in the protein databases were analyzed using GeneSilico Protein Structure Prediction meta-server (Kurowski and Bujnicki, 2003), together with PSI-BLAST (https://blast.ncbi.nlm.nih.gov/Blast.cgi) (Altschul and Koonin, 1998). The functions with the highest scores obtained from the screen were judged to be the most probable.

### Characterization of D. solani transcriptional units disrupted by the presence of Tn5 transposon

The putative transcriptional organization of *D. solani* IPO 2222 genes (single gene vs. operon) interrupted by Tn5 was established using Operon-mapper (https://biocomputo.ibt.unam.mx/operon_mapper/) (Taboada et al., 2018). The complete genome sequence of *D. solani* IPO 2222 (Genbank accession: CP015137) (Khayi et al., 2016) was used as a reference. Inference of the biochemical pathways in which the genes of interest might participate was made using KEGG (Kanehisa and Goto, 2000). The results were visualized using iPath (Letunic et al., 2008). Likewise, proteins were evaluated for their predicted biological, functional, and metabolic roles in cellular networks using STRING (Search Tool for Retrieval of Interacting Genes/Proteins) v11.5 accessed *via* the website (https://string-db.org/) (parameters: network type: *full network*, network edges: *high confidence*, interaction sources: t*ext mining, experiments, databases, co-expression, cooccurrence, gene fusion*), providing essential information regarding interactions of proteins of interest (Szklarczyk et al., 2019) using the proteome of *D. solani* strain IPO 2222 WT (van der Wolf et al., 2014) as a reference.

### Kinetics of □D5 adsorption to D. solani IPO 2222 and D. solani Tn5 mutants

The rate of □D5 adsorption to wild-type *D. solani* IPO 2222 and selected phage-resistant Tn5 mutants was determined similarly as before (Czajkowski et al., 2013a; Bartnik et al., 2021). Briefly, log-phase grown bacterial cultures were infected with a phage suspension (at Multiplicity of Infection (MOI) equal to 0.01) and incubated at 28 °C for the total time of 20 min. Two individual samples per bacterial strain to be tested were collected after 0 (control), 1, 2, 5, 10, 15, and 20 min, and assayed for free, unadsorbed phages. For negative control, bacteriophages were suspended in sterile TSB and incubated for 20 min under the same conditions as described above. The experiment was separately replicated three times with the same setup, and the obtained counts were averaged for analyses. Phage adsorption was calculated using the equation: percentage adsorption = (the average titer of unabsorbed phages per sample/ average titer of phages in negative control) x 100 (Bartnik et al., 2021).

### Kinetics of □D5 adsorption to chloramphenicol-killed D. solani IPO 2222

To test whether bacteriophage □D5 can adsorb to non□viable (dead) *D. solani* IPO 2222 cells, the dead cell adsorption assay was employed (Shao and Wang, 2008; Bartnik et al., 2021). Briefly, IPO 2222 WT was grown for 16 h in TSB at 28 °C with shaking (150 rpm). After incubation, chloramphenicol (Sigma□Aldrich, Darmstadt, Germany) was added to the final concentration of 5 mg mL^-1^ to kill bacterial cells. Such treated *D. solani* bacterial cultures were incubated under the same experimental conditions for one more hour. To test the efficiency of killing, 100 μL aliquots of bacterial culture were collected, in duplicate, plated on TSA plates, and incubated at 28 °C to allow bacterial colonies to grow. The chloramphenicol□killed *D. solani* IPO 2222 cells were mixed with a phage suspension (at MOI = 0.01) and analyzed for phage adsorption as described above. Three repetitions were done using the described procedure, and the results were averaged for analysis. Phage adsorption was calculated as described above.

### Determination of the generation time of D. solani phage-resistant Tn5 mutants in rich and minimal media

To determine whether the Tn5 insertions affect the generation time of the mutants, the growth of the selected *D. solani* Tn5 phage-resistant mutants was assessed both in TSB (rich medium) and in M9 with 0.4% glucose (minimal medium) at 28 °C for 16 h as previously described (Czajkowski et al., 2017b). The experiment was replicated one time, and the results were averaged. The average generation time of each Tn5 mutant was analyzed using the Doubling Time calculator (parameters: C0= 3 h, Ct=7 h, t=4 h) (http://www.doubling-time.com/compute.php) (Roth, 2006).

### Growth of phage-resistant Tn5 mutant at different temperatures

The growth of ΦD5-resistant Tn5 mutants and the IPO 2222 WT strain was tested on solid rich (TSA) and minimal (M9+glucose) media over a range of six temperatures: 4, 8, 15, 28, 37, and 42 °C as described before (Krzyzanowska et al., 2019). For this, 5-µl aliquots of 50-fold diluted in TSB or M9+glucose overnight bacterial cultures grown in either TSA and M9+glucose, respectively, were placed on the surface of either TSA or M9+glucose and incubated for 120 h at 5 and 8 °C or for 48 h for all other tested temperatures (15, 28, 37 and 42 °C). Growth was assessed visually daily. The experiment was replicated one time using the same setup.

### Determination of the average generation time of D. solani Tn5 mutants at low (5.0) and high (10.0) pHs

To assess whether the transposon-mediated □D5-resistance affected the generation time of the *D. solani* mutants grown at different pHs, the growth of selected *D. solani* ΦD5-resistant Tn5 mutants was measured in TSB with pH 5 and TSB with pH 10, similarly to other studies (Czajkowski et al., 2012b). Briefly, overnight cultures (density of ca. 10^9^ CFU ml^-1^ in TSB) were diluted 50-times in a fresh growth broth (TSB pH 5 or TSB pH 10). One hundred microliters of 50-times diluted bacterial culture were aseptically transferred to the wells of 96-well microtiter plates (NEST) and sealed with optically clear sealing tape (Sarstedt) to prevent evaporation. The bacterial growth rate was determined by measuring the optical density (OD) (λ=600 nm) every 0.5 h for a total time of 12 h in an Epoch2 Microplate Spectrophotometer (BioTek). The experiment was repeated once, and the obtained results were averaged. The generation time was calculated using the Doubling Time calculator (parameters: C0= 3 h, Ct=7 h, t=4 h) (http://www.doubling-time.com/compute.php) (Roth, 2006).

### Phenotypes of phage-resistant D. solani Tn5 mutants analyzed using BIOLOG Phenotypic Microarrays

Selected phage-resistant *D. solani* IPO 2222 mutants were analyzed with the BIOLOG phenotypic microarray system with GEN III and EcoPlate microplates (Biolog Inc.) as described earlier (Bartnik et al., 2021). Prior to BIOLOG analysis, bacterial cultures were grown on TSA plates for 24 h at 28°C, and then carefully resuspended into inoculation fluid (IF-A) (GENIII) or into 10 mM phosphate buffer pH PBS ma 7.4 (EcoPlate) using a sterile cotton swab (Bionovo). The turbidity of the bacterial suspensions was adjusted to ca. 90 % T with a spectrophotometer [A = log(%T)]. One hundred microliters of bacterial suspensions in duplicates were inoculated into each well of the 96-well microplates. Inoculated 96-well plates were sealed with optically clear sealing tape (Sarstedt) and incubated for 24 h at 28°C. The wells were then examined for a colour change (appearance of colour = positive reaction). Colour development was also documented using an Epoch2 microplate spectrophotometer (BioTek) equipped with a λ = 595-nm wavelength filter. Plates inoculated with the wild-type *D. solani* strain IPO 2222 WT were used as controls.

### Phenotypes of phage-resistant D. solani Tn5 mutants analyzed using plate assays

□D5-resistant *D. solani* Tn5 mutants were screened for phenotypic features, putatively important for their interaction with plant tissues and/or environmental fitness, including swimming and swarming motility (Czajkowski et al., 2012b), biofilm formation (Shao et al., 2019), the ability to grow on the TSA medium supplemented with 5% NaCl (Dickey, 1979), ability to grow in TSB at different pHs (Czajkowski et al., 2012b), production of enzymes: pectinolytic enzymes (Perombelon and van Der Wolf, 2002), cellulases (Py et al., 1991), proteases (Ji et al., 1987) and siderophores (Schwyn and Neilands, 1987).

### Phenotypes of phage-resistant D. solani Tn5 mutants analyzed using whole□cell MALDI-TOF-MS Analyses

*D. solani* IPO 2222 Tn5 phage□resistant mutants were tested using a whole□cell MALDI□TOF MS spectral analysis as previously described (Bartnik et al., 2021). Briefly, IPO 2222 WT and Tn5 mutants were grown on TSA at 28 °C for 24 h. Viable cells were loaded with a sterile inoculation loop directly from the culture onto a MALDI plate. Mixture 1:1 of ferulic acid (FA, 10 mg/ml in 33% acetonitrile, 13% formic acid, water) and dihydroxybenzoic acid (DHB, 10 mg/ml in 50% acetonitrile in water and 0.1% trifluoroacetic acid) was used as a matrix. 1 μL of matrix solution was used to overlay the spot containing bacterial samples, and such prepared plate was left to crystallize at room temperature. After preparation of the spots (ca. 15 min), protein mass fingerprints were obtained using a 5800 MALDI-TOF/TOF mass spectrometer (AB Sciex, Framingham, MA, USA), with detection in the linear middle mass in the range from 5000 to 20 000 Da, positive ion mode for a total of 1000 laser shots with a 1 kHz OptiBeam laser (YAG, 349 nm) (Bartnik et al., 2021). Laser intensity was corrected for all tested samples. Registered MS spectra were examined with Data Explorer software (AB Sciex). All MALDI□TOF MS spectra reported in this study were averages of six replicated measurements (2 independent measurements, each containing 3 technical repetitions) per analyzed strain as described before (Bartnik et al., 2021).

### Cell and colony morphology of phage-resistant Tn5 D. solani mutants analyzed with light and transmission electron microscopy

The colony morphology of selected phage-resistant *D. solani* IPO 2222 Tn5 mutants was analyzed using a Leica MZ10 F stereomicroscope with 10× and 40× magnifications combined with a Leica DFC450C camera (Leica) as previously described (Lisicka et al., 2018; Bartnik et al., 2021).

The morphology of phage resistant Tn5 *D. solani* mutants was evaluated using transmission electron microscopy (TEM) as previously described (Czajkowski et al., 2017b). TEM analyses were performed at the Laboratory of Electron Microscopy (Faculty of Biology, University of Gdansk, Poland). For that, bacteria were adsorbed onto carbon-coated grids (GF Microsystems), directly stained with 1.5% uranyl acetate (Sigma-Aldrich), following their analyses with an electron microscope (Tecnai Spirit BioTWIN, FEI) as described previously (Czajkowski et al., 2017b). At least ten images of each mutant and the wild-type strain were taken to estimate cell diameter.

#### Cell morphology of phage-resistant Tn5 *D. solani* mutants analyzed with scanning electron microscopy (SEM)

*D. solani IPO2222* WT and phage-resistant Tn5 mutant cells were fixed for microscopic analyses according to a standard SEM procedure (Fiolka et al., 2015). Briefly, bacterial cells were fixed with 4% glutaraldehyde in 0.1 M phosphate buffer, pH 7.0. Bacterial cells were then treated with OsO4, dehydrated in a graded series of acetone concentrations (15%, 30%, 50%, 70% and twice 100%). This process was followed by drying the specimens using silica gel for 24 hours, and then the samples were gold-sputtered using a K550X sputter coater (Quorum Technologies). Finally, as previously described, cells were visualized with a Vega 3 Scanning Electron Microscope (Tescan) (Fiolka et al., 2015).

#### Cell surface morphology of phage-resistant Tn5 D. solani mutants analyzed with atomic force microscopy (AFM)

Imaging the cell surface of *D. solani* IPO2222 WT and selected phage-resistant mutants M399 and M1004 with atomic force microscopy (AFM) was performed as previously described (Fiolka et al., 2015). The measurements were taken in Peak-Force Quantitative Nanomechanical Mapping mode using NanoScope V AFM (Bruker, Vecco Instruments Inc., Billerica, MA, USA) equipped with NanoScope 8.15 software. The force constant of the NSG01 probe (NT-MDT Spectrum Instruments, Russia) was in the range 1.45 – 15.1 N/m. Three fields of 3 µm × 3 µm were scanned for each sample. The roughness was read from 10 areas of 100 nm × 100 nm from each image, and the arithmetic mean was calculated from the obtained results. Images were analyzed with Nanoscope Analysis version 1.4.

### Autoaggregation (sedimentation) of Tn5 mutants

In bacterial cultures, autoaggregation of cells leads to accelerated sedimentation and phase separation of the cultures, resulting in the bacterial cells collecting at the bottom of the culture vessel, leaving a clear supernatant above (Sorroche et al., 2010). The autoaggregation assay was performed to test whether phage-resistant Tn5 D. solani mutants are altered in aggregation. For that, the phage-resistant Tn5 bacterial mutants were assessed for the speed of autoaggregation (sedimentation) as described in (Dorken et al., 2012; Trunk et al., 2018). Briefly, IPO 2222 WT (control) and 13 phage-resistant Tn5 mutants were grown in 10 ml TSB at 28 °C for 24 h with shaking. After incubation, bacterial cultures (1 mL) were separately transferred to sterile cuvettes (Eppendorf), and the optical density (OD600) of each bacterial culture was measured (time=0 h). Next, the cuvettes were covered with parafilm and incubated for the next 24 h at 28 °C without shaking. After this time, the optic density (OD600) of each bacterial culture was measured again (time=24 h). Two replicates were done per each analyzed bacterial strain and the entire experiment was repeated one time with the same setup. The results from both repetitions were averaged for analysis. Percentage aggregation (sedimentation) was measured as follow: %A = 1-(OD60024h/OD600 0h), where: %A – percentage of aggregation (sedimentation), OD6000h – OD of bacterial culture at time 0 h, OD60024h – OD of bacterial culture at time 24 h (Dorken et al., 2012).

### Permeability of Tn5 mutants cell outer membrane

Permeability of the bacterial outer membrane was assessed by a degree of cell lysis observed during the incubation of bacteria in the presence of sodium dodecyl sulphate (SDS) (Przepiora et al., 2022). Briefly, *D. solani* WT and *D. solani* phage-resistant Tn5 mutants were grown in TSB (WT strain) or in TSB supplemented with 50 μg mL^-1^ neomycin (Tn5 mutants) for 16 h at 28 °C with shaking. After incubation, 2 ml of each bacterial culture was separately collected, washed two times with PBS pH 7.2 (Sigma-Aldrich) and resuspended in 2 ml of PBS. Optical density (OD600 = 0.1) was used to normalize the number of bacterial cells in all treatments (ca. 10^8^ CFU mL^-1^). The 180 µl aliquots of the normalized bacterial suspensions in PBS were separately transferred to the wells of 96-well plate (Greiner Bio-One). To each well containing bacterial suspension, either 20 µl of sterile water (control) or 20 µl of 10% SDS (Sigma-Aldrich) in sterile water (treatment) was added. The cell lysis was monitored at room temperature *via* a decrease of optical density (OD) of the tested bacterial culture measured every minute for the total time of 10 min using Epoch2 Microplate Spectrophotometer (BioTek). Two replicates were done per each analyzed bacterial strain and the entire experiment was repeated one time with the same setup. The results from both repetitions were averaged for analysis.

### Antibiotic susceptibility of Tn5 mutants

The antibiotic susceptibility of *D. solani* Tn5 mutants was determined by a disc diffusion method as previously described (Bauer et al., 1966; Bartnik et al., 2021). Antibiotic discs (BD BBL - Sensi-Disc antimicrobial test discs) used in this study were: chloramphenicol (30 µg), gentamicin (10 µg), tigecycline (15 µg), doxycycline (30 µg), sulfametoxazol/trimetropin (23,75/1.25 µg), ciprofloxacin (5 µg), ceftaroline (5 µg), imipenem (10 µg), piperacillin/tazobactam (30/6 µg), cefuroksym/ceftaroline (30/5 µg), cefuroxime (30 µg), aztreonam (30 µg), ampicillin (10 µg), ampicillin/sulbactam (10/10 µg), colistin (10 µg), fosfomycin (200 µg). Briefly, the wild-type *D. solani* and phage-resistant *D. solani* Tn5 mutants were grown for 16 h in TSB supplemented with neomycin (50 μg mL^-1^), and WT was grown in TSB at 28 °C with shaking (120 rpm). The Mueller-Hinton (MH medium, BD) supplemented with 1.5 % agar (Oxoid) square plates (100 x 100 mm) were inoculated using a sterile cotton swab (Sarstedt) soaked in a suspension of individual strains (WT or Tn5 mutant). Once the inoculated Mueller-Hinton plates had dried, the antibiotic discs were placed on the agar surface in such a way to ensure a minimum distance between each disc of ca. 1.5 -2 cm. Plates were incubated at 28 °C for 24 h, and afterwards, they were examined for the presence/absence of a clear halo in the bacterial lawn around the discs. The presence of the halo around the antibiotic disk was recorded as a negative reaction (bacterial susceptibility). In contrast, the lack of halo was registered as a positive reaction (bacterial resistance). *D. solani* strain IPO 2222 WT was used as a control. The experiment was repeated once.

### Isolation and visualization of lipopolysaccharide (LPS) from wild-type D. solani strain IPO 2222 and phage-resistant D. solani Tn5 mutants

Lipopolysaccharides (LPS) of *D. solani* IPO 2222 WT strain and ΦD5□resistant Tn5 mutants were isolated using a Lipopolysaccharide Extraction Kit (Abcam, Symbios, Gdansk, Poland) with a modified protocol (Bartnik et al., 2021). Resulting lipopolysaccharides were separated using gradient 4–20% sodium dodecyl sulfate□polyacrylamide gel (4–20% Mini□PROTEAN® TGX™ Precast Protein Gel, BioRad, Hercules, USA) electrophoresis (SDS□PAGE) according to (Sambrook et al., 1989) and visualized with silver staining as described before (Tsai and Frasch, 1982).

### Population dynamics of D. solani phage-resistant mutants on S. tuberosum adaxial leaf surface in vitro

*S. tuberosum* potato plants, cultivated as described earlier (Czajkowski et al., 2017a), were grown in a phytochamber at 22°C and 16/8 h (day/night) photoperiod (white cool fluorescent light, Philips, TLD 58 W/84o, 30-35 μmol m^-2^ s^-1^) and 80% relative humidity (RH). *S. tuberosum* leaves (third to sixth from the shoot tip) were detached from the plants and used for detached leaf assay experiment as described earlier (Fikowicz-Krosko and Czajkowski, 2018). Briefly, two ml of bacterial suspension (IPO 2222 WT or selected □D5-resistant IPO 2222 Tn5 mutants) adjusted to 10^6^ CFU mL^-1^ in sterile Ringer’s buffer (Merck) was sprayed using a manual sprayer on the adaxial surface of each detached leaf. Sprayed leaves were dried in the laminar flow and placed, adaxial side up, on 0.5% water agar (Oxoid) supplemented with 200 µg mL^-1^ cycloheximide (Sigma-Aldrich), to prevent fungal growth, in square plastic Petri dishes (100x100 mm, Sarstedt). Sterile Ringer’s buffer was used instead of bacterial suspensions as a control. The inoculated leaves were kept in the growth chamber under the same conditions as described above. Samples were collected at 0 and 14 days post inoculation (dpi) and assayed for the presence of *D. solani* IPO 2222 WT strain or □D5-resistant Tn5 mutants. At each time point, per treatment, 5 randomly chosen leaves were collected and separately shaken at 50 rpm in 10 ml of Ringer’s buffer volume at room temperature in 50 ml Falcon tubes (Sarstedt), each for 30 min to wash out bacterial cells from the surfaces of the leaves. The washings containing bacterial cells were serially diluted in Ringer’s buffer and 100 μl taken from the 10 and 100 times diluted samples were plated in duplicates on CVP (for IPO 2222 WT) supplemented with 200 μg ml^-1^ cycloheximide (Sigma) or on CVP containing 50 μg ml^-1^ neomycin and 200 μg ml^-1^ cycloheximide to select for phage resistant Tn5 mutant colonies. The inoculated plates were incubated at 28°C, and the neomycin-resistant, cavity-forming *D. solani* colonies were counted. The experiment was replicated one time with the same setup, and the results were averaged.

### The ability of selected phage-resistant mutants to cause maceration of potato tubers

Potato tubers of cv. Bryza were obtained locally in Gdansk, Poland. Five individual potato tubers were inoculated with a given mutant and assessed for infection symptoms, using a whole tuber injection method (stab inoculation), as described before (Czajkowski et al., 2012b; Czajkowski et al., 2017b).

For this, per tuber, a volume of 100 µl of bacterial culture was delivered to potato tuber by stab inoculation into tuber pith using a 200 µl-volume yellow tip filled with 100 µl bacterial suspension. Wild□type *D. solani* IPO 2222 was used as a positive control, and the negative control was sterile demineralized water. Five replications were made for each tested mutant (five individual tubers, n=5), and the experiment was one time replicated with the same setup (n=10).

### Virulence of D. solani Tn5 mutants in potato plants

Replicated experiments of plants grown in a growth chamber were performed in November and December 2021 using the previously developed protocol (Czajkowski et al., 2017a). The certified potato tubers of cv. Kondor were acquired from Plant Breeding and Acclimatization Institute – National Research Institute, Bonin, Poland. *S. tuberosum* potato plants were cultivated as described earlier (Czajkowski et al., 2017a). After two weeks of cultivation, rooted plants with a height of ca. 10-15 cm were transferred to 1 L pots and cultivated in potting compost for the next 2 weeks under similar growth conditions. One h before the infestation of soil with bacteria, plants were watered to keep the soil moist during the infestation. Potato plants (n= 5 plants per repetition, 10 per treatment) were infected with bacterial strains by the application of bacterial suspensions (50 ml of 10^8^ CFU mL^-1^ bacterial suspension in sterile 1/4 Ringer’s buffer) directly to the soil surrounding stems. As a negative control, plants were watered with sterile Ringer’s buffer (50 ml per plant) instead of bacterial suspensions. Pots were randomized for the growth chamber experiments: 5 blocks of 15 pots per treatment (n=75 plants per repetition), and the experiment was one time replicated using the same setup (total n=150 plants). Plants were visually inspected daily for the development of the disease symptoms: chlorosis, black rotting of the stem, haulm wilting, and the death of the plants. Plants were sampled at 14 dpi (days post-inoculation) by cutting ca. 2 cm long stem segments, located ca. 5 cm above ground level, and pooled per analyzed plant (Bartnik et al., 2021). The stem fragments were surface-sterilized as described before (Czajkowski et al., 2017a). The presence of *D. solani* cells inside potato stems was determined by plating the stem extracts on CVP (*D. solani* IPO 2222 WT) supplemented with 200 μg ml^-1^ cycloheximide and/or CVP supplemented with 50 μg mL^-1^ neomycin and 200 μg ml^-1^ cycloheximide (□D5-resistant *D. solani* Tn5 mutants) and counting the resulting bacterial colonies. The results were averaged for analysis.

### Competition of wild type IPO 2222 and selected phage-resistant Tn5 mutants on potato tubers

*Generation of fluorescently-tagged D. solani and D. solani phage-resistant Tn5 mutants* Plasmids pPROBE-AT-*gfp* (Miller et al., 2000) and pRZ*-*T3*-dsred* (Bloemberg et al., 2000), both used previously for stable tagging of *D. solani* cells with fluorescent proteins (Czajkowski et al., 2010; Czajkowski et al., 2013b), were used to generate the GFP-tagged and dsRed-tagged variants of phage-resistant *D. solani* Tn5 mutants and/or IPO 2222 WT, respectively. The pPROBE-AT-*gfp* and pRZ*-*T3*-dsred* plasmids were introduced into bacterial cells by electroporation as described in (Czajkowski et al., 2010). After electroporation, bacterial cultures (100 µL) were plated on TSA plates containing 100 µg mL^-1^ ampicillin (for selection of GFP-positive colonies) or on TSA plates containing 40 µg mL^-1^ tetracycline (for selection of dsRed-positive colonies) and incubated for 24-48 h at 28 °C to allow bacteria to grow. The obtained GFP and dsRed tagged Tn5 strains were routinely grown on TSA supplemented with neomycin (50 µg mL^-1^) and additionally supplemented with ampicillin (100 µg mL^-1^) for selection of GFP-tagged strains or tetracycline (40 µg mL^-1^) for selection of DsRed-tagged *D. solani* strains unless stated otherwise. In addition, the growth of fluorescent protein-tagged strains was evaluated as previously described (Czajkowski et al., 2010).

### Co-inoculation of potato tubers with fluorescently tagged D. solani WT and Tn5 ΦD5-resistant mutants and sampling of tubers for PT-pour plating

To determine whether phage-resistant mutants may be altered in their fitness during infection of potato tubers compared to the IPO 2222 WT, two independent tuber co-inoculation experiments were conducted in November 2021. In the first experiment, a GFP-tagged *D. solani* IPO 2222 WT (IPO 2254) (Czajkowski et al., 2010) strain together with DsRed-tagged phage-resistant Tn5 *D. solani* mutants were used, whereas in the second experiment, DsRed-tagged IPO 2222 WT (IPO 2222-DsRed) and GFP-tagged phage-resistant mutants Tn5 mutants were applied. The fluorescently tagged *D. solani* IPO 2222 (either IPO2254 or IPO 2222-DsRed) was grown in TSB supplemented with ampicillin (100 µg mL^-1^) or tetracycline (40 µg mL^-1^), respectively for 16 h at 28 °C with shaking (200 rpm). Fluorescently tagged *D. solani* phage-resistant Tn5 mutants (tagged either with GFP or DsRed) were grown under the same conditions, but the bacterial growth medium was additionally supplemented with neomycin (Sigma-Aldrich) to a final concentration of 50 µg mL^-1^. After incubation, 5 mL of bacterial cultures were separately collected, centrifuged (6000 x RCF, 5 min.), washed twice with sterile 1/4 Ringer’s buffer and finally resuspended in initial volume (5 mL) of sterile Ringer’s buffer. Bacterial suspensions were diluted in sterile Ringer’s buffer to a density of ca. 10^8^ CFU mL^-1^ (optical density at 600 nm [OD600] = 0.1). Potato tubers of cultivar Bryza (n=5 per mutant, per repetition, n=10 per experiment) were obtained locally in Gdansk, Poland. Prior to other experiments, 25 potato tubers, randomly selected from the pool of the tubers used in the experiment, were tested for the absence of *Pectobacterium* spp. and *Dickeya* spp. as described earlier (Czajkowski et al., 2009b). *Pectobacterium* spp. and *Dickeya* spp.-free tubers were rinsed with running tap water to remove soil particles, surface-sterilized for 20 min in 5% commercial bleach solution in water, and washed twice for 1 min with demineralized, sterile water. Surface-sterilized tubers were dried under laminar flow. Inoculation of tubers with bacteria was done using a whole tuber injection method as described earlier (Czajkowski et al., 2012b; Czajkowski et al., 2017b). For this, per tuber, a volume of 100 µl of bacterial culture was delivered to potato tuber by stab inoculation into tuber pith using a 200 µl-volume yellow tip filled with 100 µl bacterial suspension. Treatments contained either (per 100 µl): (i): 10^6^ CFU mL^-1^ of *D. solani* IPO 2222 WT strain, (ii): 5 x 10^5^ CFU mL^-1^ of *D. solani* IPO 2222 together with 5 x 10^5^ CFU mL^-1^ of individual phage-resistant *D. solani* Tn5 mutant (13 individual treatments) or (iii): 10^6^ CFU mL^-1^ of phage-resistant *D. solani* Tn5 mutant alone. Inoculated tubers were kept in humid boxes (80-90% relative humidity) at 28 °C for 72 h to allow bacteria to rot potato tissue. After incubation, per tuber, ca.

1.5 -2 g of rotten potato tissue was collected and resuspended in the volume of Ringer’s buffer to twice the weight of the sample (Czajkowski et al., 2009b). To protect bacterial cells from oxidative stress, the Ringer’s buffer was supplemented with an oxidant, 0.02% diethyldithiocarbamic acid (DIECA, Sigma-Aldrich). One hundred µl of undiluted, 10-fold- and 100-fold diluted samples, in duplicates, were mixed with 300 µL of PT medium (Perombelon and van Der Wolf, 2002) prewarmed to 45-50 °C and containing 200 µg mL^-1^ cycloheximide, 50 µg mL^-1^ neomycin and either 150 µg mL^-1^ ampicillin (for selection ofthe GFP-tagged strains) or 40 µgmL^-1^tetracycline (for selection of the DsRed-tagged strains) in the well of a 48-well plate (Greiner Bio-One) . Plates containing solidified medium were incubated for 24-48 h at 28 °C to allow bacteria to grow. Wells were inspected for the presence of GFP-and DsRed-fluorescent bacterial cells using an epifluorescence stereomicroscope (Leica MZ10 F and Leica DFC450C camera system) (Czajkowski et al., 2012a). The GFP-positive and DsRed-positive colonies were counted. The experiment was replicated one time with the same setup. The results from both repetitions were averaged for analysis.

### Statistical analyses

Statistical analyzes were performed as previously described (Bartnik et al., 2021). Briefly, bacterial colony counts were transformed as log(x + 1). The Shapiro–Wilk test (p <0.05) (Shapiro and Wilk, 1965) was used to test whether the distribution of the results for individual counts follows a normal distribution, whereas, for the populations of counts that were not equally scattered between the analyzed groups (not normally distributed) (e.g. control vs. treatment or treatment vs. treatment), the Welch’s T□test was applied (Welch, 1947). The variance homogeneity was validated using the Fisher– Snedecor test (Box, 1953). Pair□wise differences were evaluated using a two□tailed Student’s t□test (Student, 1908). The treatments involved potato tubers, plants in growth chamber and potato leaves were analyzed, matching the experimental pattern in which two replicated experiments were done per each treatment of replicated plants/tubers/leaves. The linear model was a complete block design with replicates as individual blocks (Shieh and Jan, 2004). The main effects observed were analyzed for the impact of time and treatment type and a two□way interaction between time and treatment type.

## Results

### Transposon mutagenesis and identification of Tn5-disrupted genes in ΦD5-resistant *D. solani* IPO 2222 mutants

The 1000 *D. solani* Tn5 mutants were randomly selected from the pool of 10000 mutants obtained in our former studies (Lisicka et al., 2018; Czajkowski et al., 2020) and screened for the resistance to infection caused by bacteriophage vB_Dsol_D5 (ΦD5). A total of 13 (ca. 1.3%) *D. solani* Tn5 mutants were found to be resistant to phage ΦD5 (Tab. 1). The genomes of these 13 phage-resistant *D. solani* mutants were sequenced to identify the Tn5 insertion sites. A single insertion of the transposon Tn5 was found in each of the 13 ΦD5-resistant *D. solani* mutants analyzed in this study (data not shown).

**Table 1.**
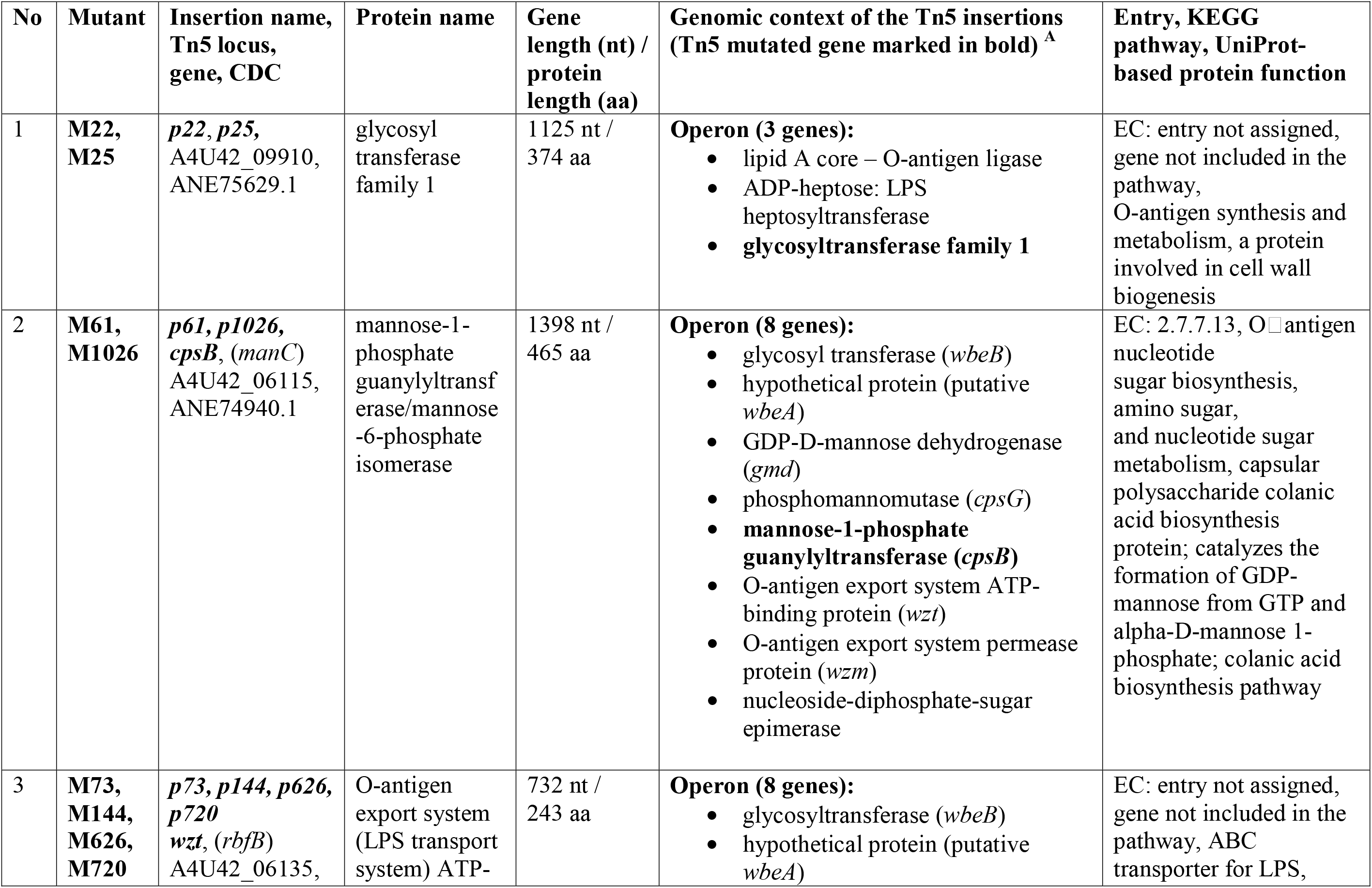
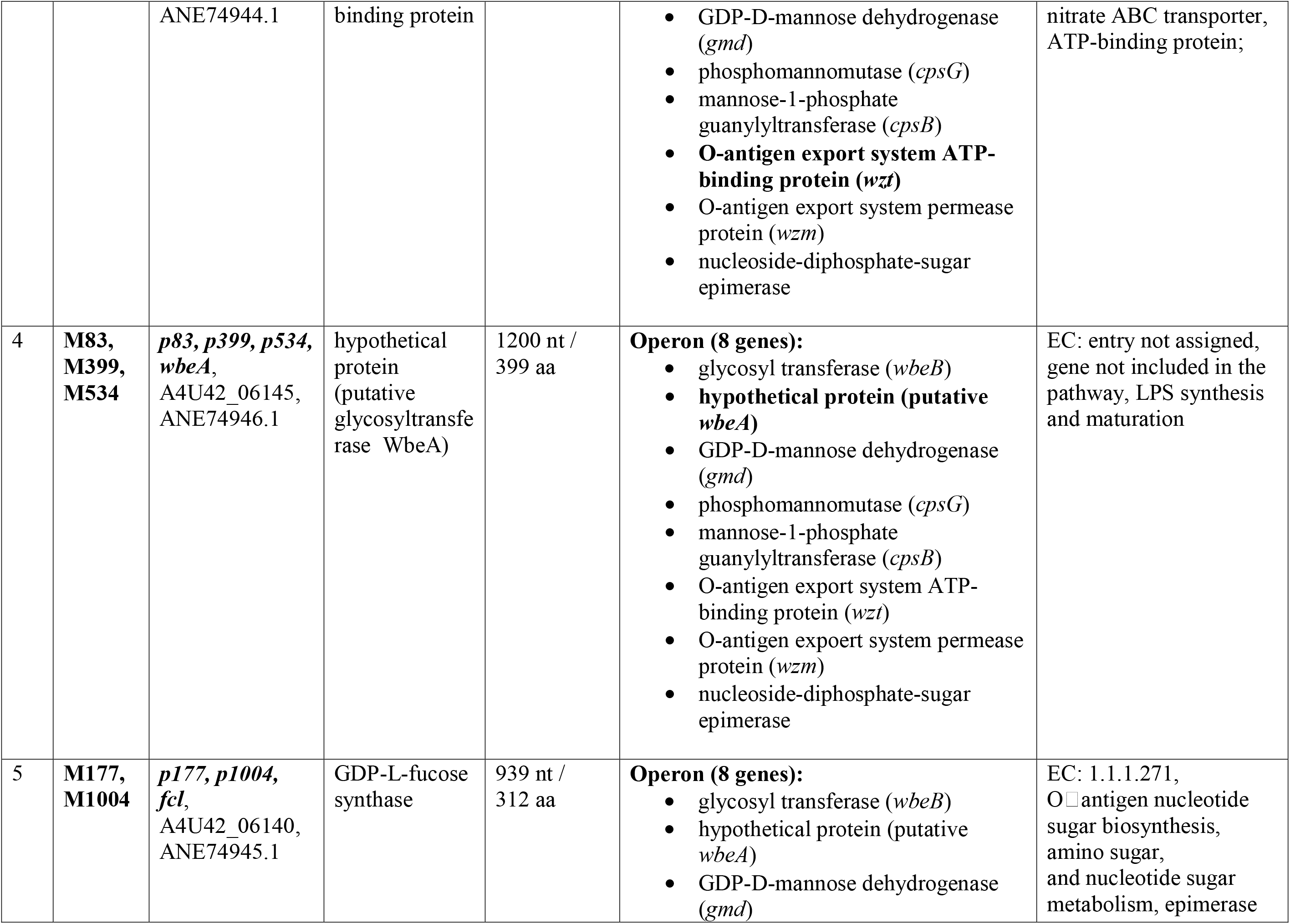
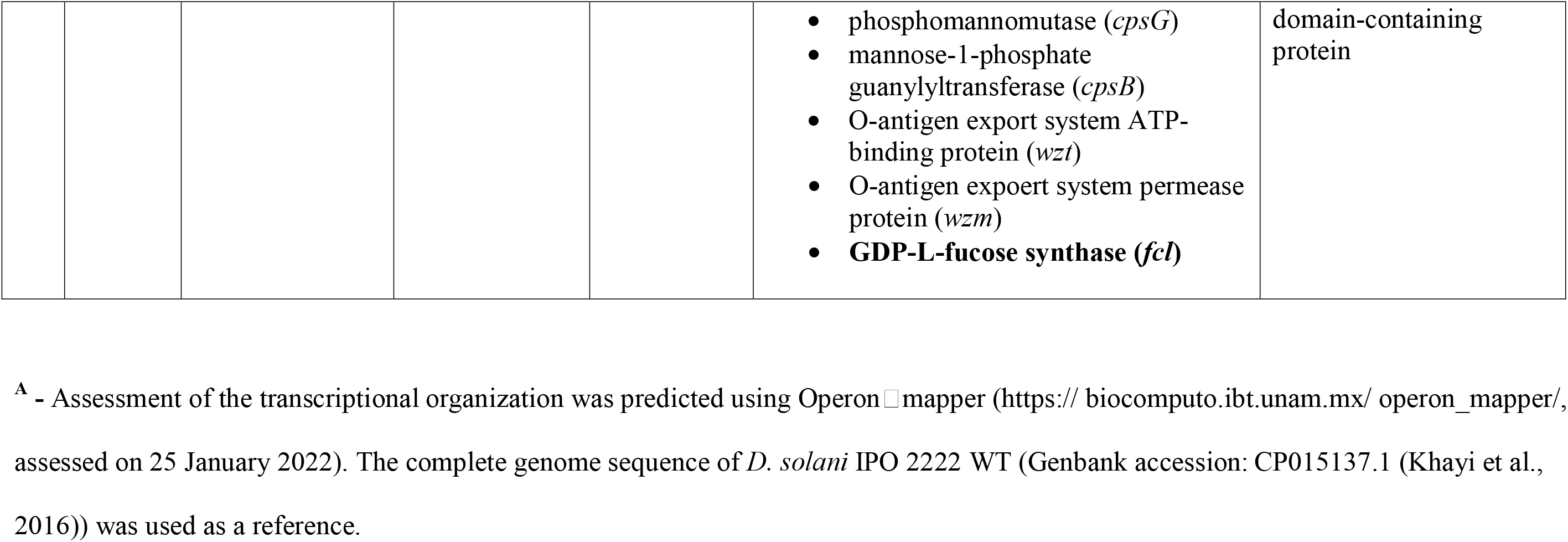
Genetic loci of *Dickeya solani* strain IPO 2222 Tn5 mutants expressing resistance against phage vB_Dsol_D5 (ΦD5)

### Characterization of the Tn5-disrupted genes in phage-resistant *D. solani* mutants

All 13 loci disrupted by the presence of Tn5 transposon in ΦD5-resistant *D. solani* IPO 2222 mutants encoded proteins associated with synthesis, metabolism, storage and/or modification of bacterial surface including LPS, EPS and capsule (CPS) (Table 1, Fi.g 1). Interestingly, multiple times the Tn5 transposon targeted the same locus in the case of mutant M22 and M25 (locus: glycosyltransferase family 1), mutant 61 and M1026 (locus: mannose-1-phosphate guanylyl-transferase/ mannose-6-phosphate isomerase), mutants M73, M144, M626 and M720 (locus: sugar ABC transporter ATP-binding protein), mutants M83, M399 and M534 (locus: hypothetical protein homological to protein WbeA – involved in O-antigen transport system) and mutants M177 and M1004 (locus: GDP-fucose synthase) (Table 1, Fig. 1). In total, five distinct bacterial loci were found to be involved in the interaction of *D. solani* IPO 2222 with bacteriophage ΦD5. The sequences bordering these five loci (ca. 1000-5000 bp.) were analyzed using BlastP to obtain additional insights into their genomic context. Of the five individual genome regions that were interrogated for their transcriptional organization, all five insertions were predicted to be parts of only 2 operons (Fig. 1, Table 1): (i) operon, which is a part of the *rfa* gene cluster involved in the biosynthesis of the core region of LPS (insertions in M22 and M25 mutants) and (ii) putative O-antigen LPS biosynthesis cluster (Ranjan et al., 2021) (insertions in M61, M73, M83, M144, M177, M399, M534, M626, M720, M1004 and M1026 mutants). Examination of the KEGG biochemical pathways corresponding to these 5 transcriptional units enabled their assignment to the cellular pathways involved in the biosynthesis of cell surface lipopolysaccharides and exopolysaccharides. Likewise, the putative interacting partners of the 5 *D. solani* proteins were assessed using STRING. STRING analyses revealed that all 5 protein-coding loci interact with proteins associated with bacterial cell surface (synthesis and remodelling of the capsule, LPS and EPS) (Supplementary Table 2). The five loci found in this study are all conserved in other members of the *Dickeya* genus; their homologs were found in *D. dianthicola*, *D. dadantii*, *D. fangzhongdai*, *D. zeae*, *D. oryzae*, *D. unidicola* and *D. chrysanthemi* strains (Supplementary Table 2).

**Figure 1.**
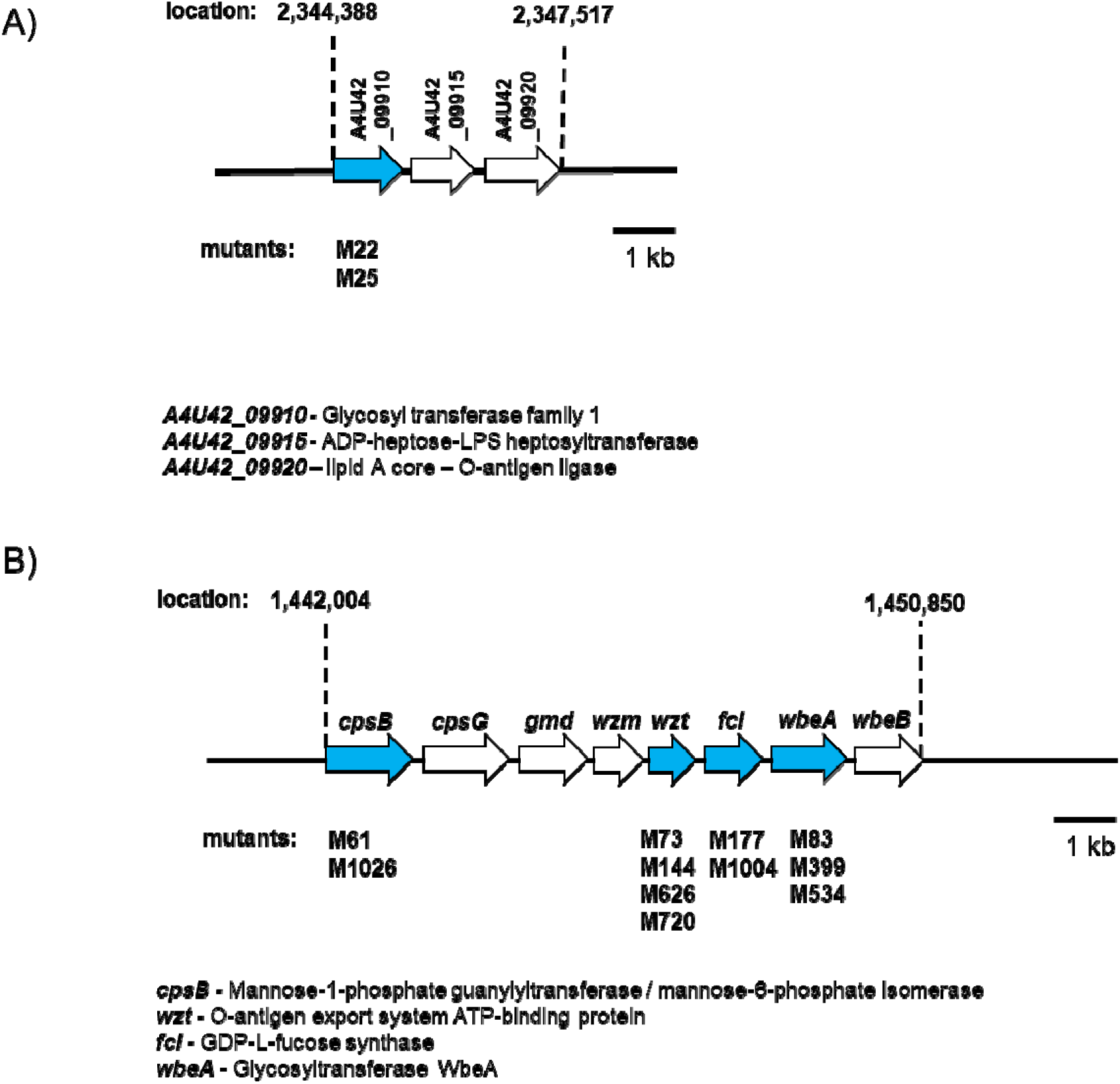
Two operons of *D. solani* strain IPO 2222 involved in the interaction of the bacterium m with lytic bacteriophage ΦD5. (A) Operon associated with *rfa* gene cluster involved in thee biosynthesis of the core region of LPS in Gram-negative bacteria and (B) Putative O-antigen LPS biosynthesis cluster (Ranjan et al., 2021). The *D. solani* IPO 2222 ORFs affectctedby the Tn5 insertion are marked in blue. The directions of the arrows represent the direction of of the transcription. The complete genome sequence of *D. solani* strain IPO2222 (GenBank accession: CP015137.1) was used to visualize the genetic organization of the clusters.

### Adsorption of ΦD5 to *D. solani* WT and *D. solani* mutant cells *in vitro*

The adsorption of the vB_Dsol_D5 (ΦD5) to the viable and non-viable (chloramphenicol-killed) cells of wild-type *D. solani* IPO2222 cells was rapid (Fig. 1). Within the first 5 min., ca. 90% of phage particles has adsorbed to the viable host cells, whereas in the total assay time (20 min.), more than 99% of ΦD5 particles have adsorbed to the host IPO 2222 WT cells under experimental conditions. Furthermore, the rate of adsorption of the ΦD5 to IPO 2222 WT cells killed by antibiotic prior to the adsorption experiments was similar to that of the viable WT cells. In contrast, the adsorption of ΦD5 to 11 phage-resistant Tn5 mutants (M22, M25, M61, M73, M83, M144, M399, M534, M626, M720, M1026) was entirely abolished, whereas adsorption to two other mutants (M177 and M1004) was significantly reduced (Fig. 1). In M177 and M1004 mutants, only between 1 and 7% of the phage particles had adsorbed to the surface of IPO2222 cells by 20 min., with the rest remaining free (Fig. 1). The results of adsorption experiments were confirmed using transmission electron microscopy (TEM), which was used to visually assess the interaction of phage ΦD5 with IPO 2222 WT and with the 13 phage resistant Tn5 mutants. In repeatable experiments, the abundant adsorption of the phage particles to the surface of IPO 2222 WT cells was observed, whereas no adsorption of ΦD5 was detected in the case of the phage-resistant Tn5 mutants tested (Fig. 2).

**Figure 2.**
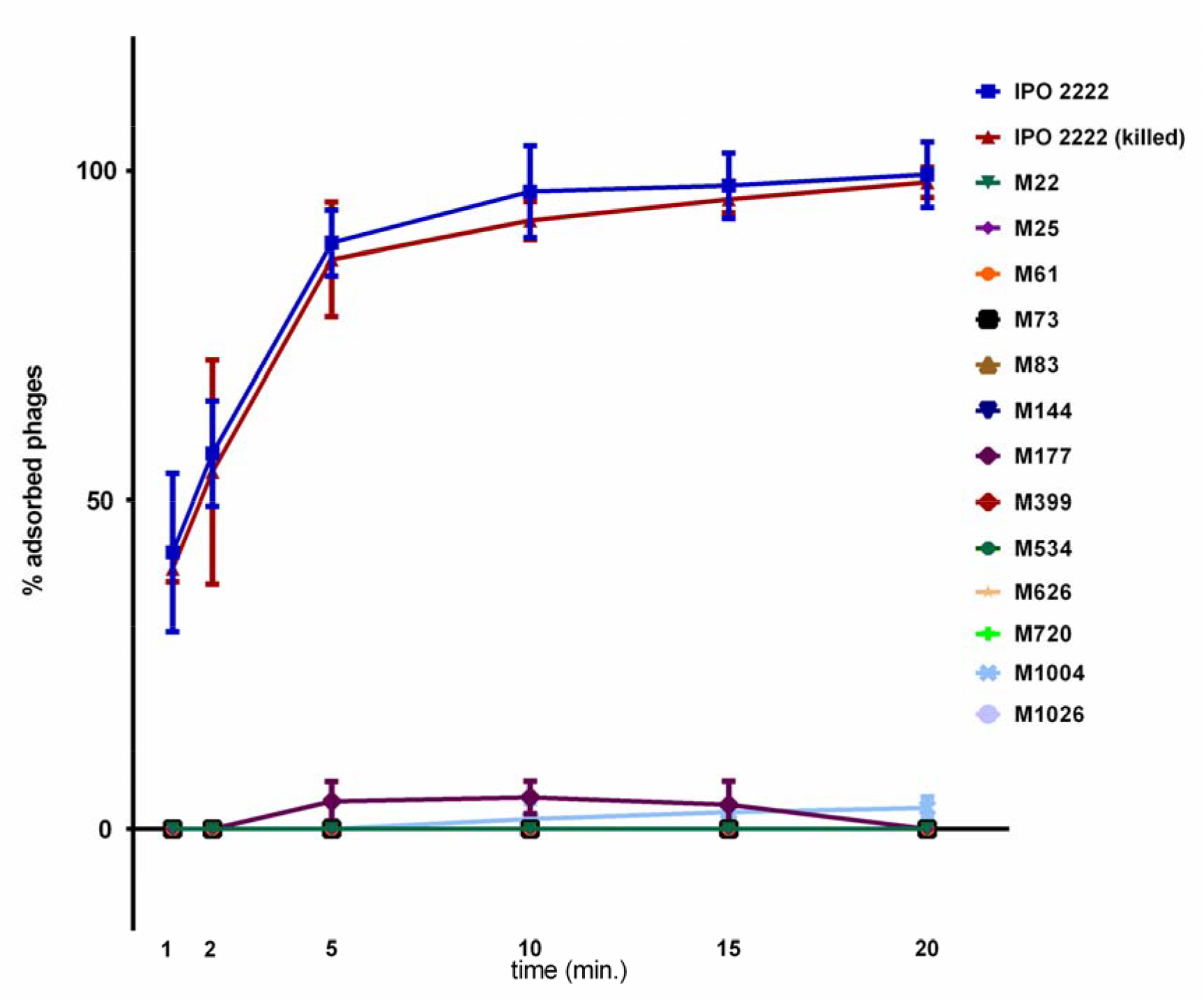
Adsorption of vB_Dsol_D5 (ΦD5) to viable D. solani IPO 2222 WT, IPO 2222 WT killed with chloramphenicol, and 13 phage-resistant D. solani Tn5 mutants. A MOI of 0.01 of D5 was used for adsorption assay and the total assay time was 20 min. Phage adsorption was calculated as follows: the percentage adsorption = (the average titer of unabsorbed phages per sample/average titer of phages in negative control) ×100. The averages and standard deviations of three independent repetitions per strain (WT or mutants) are shown.

### Analysis of the permeability of *D. solani* and Tn5 mutants outer membrane

As the mutations in all ΦD5-resistant *D. solani* mutants suggested that phage resistance is related to the alterations of bacterial envelope components (e.g. capsule, LPS, EPS, cell surface), the WT strain and 13 phage-resistant mutants were assessed for permeability of their cell membranes. In repeatable experiments, a 30% reduction (70% survival) of IPO 2222 WT (positive control) cell numbers was observed already after 1 min. incubation with SDS, whereas only ca. 15% of WT cells survived 10 min. incubation of bacterial culture in the presence of SDS. The decline of the cell numbers in the presence of SDS was significantly speeded up in the case of all phage-resistant Tn5 mutants tested, in which an average survival of cells in the presence of SDS after 1 min. assay time was between only 30 and 60% (reduction of cell numbers of 40-70%) depending on the mutant tested.

### Characterization of the lipopolysaccharide (LPS) isolated from *D. solani* IPO 2222 WT and Tn5 mutants

To evaluate whether LPS is involved in ΦD5 adsorption to IPO 2222 cells, preparations of the crude LPS from WT strain IPO 2222 and the 13 Tn5 mutants were compared using SDS-PAGE coupled with silver staining. The lipopolysaccharides from all tested strains were resolved into several bands characterized by different mobilities (Fig. 3). The presumed lipid A-core of the LPS derived from the WT strain was composed of two bands (one heavy and one faint) with masses less than 11 kDa. The putative O-antigen component of the IPO 2222 WT LPS was composed of 7 bands with masses between 11 and 48 kDa. All LPSs purified from the ΦD5-resistant *D. solani* Tn5 mutants resembled that of the wild-type IPO 2222 strain (Fig. 3).

**Figure 3.**
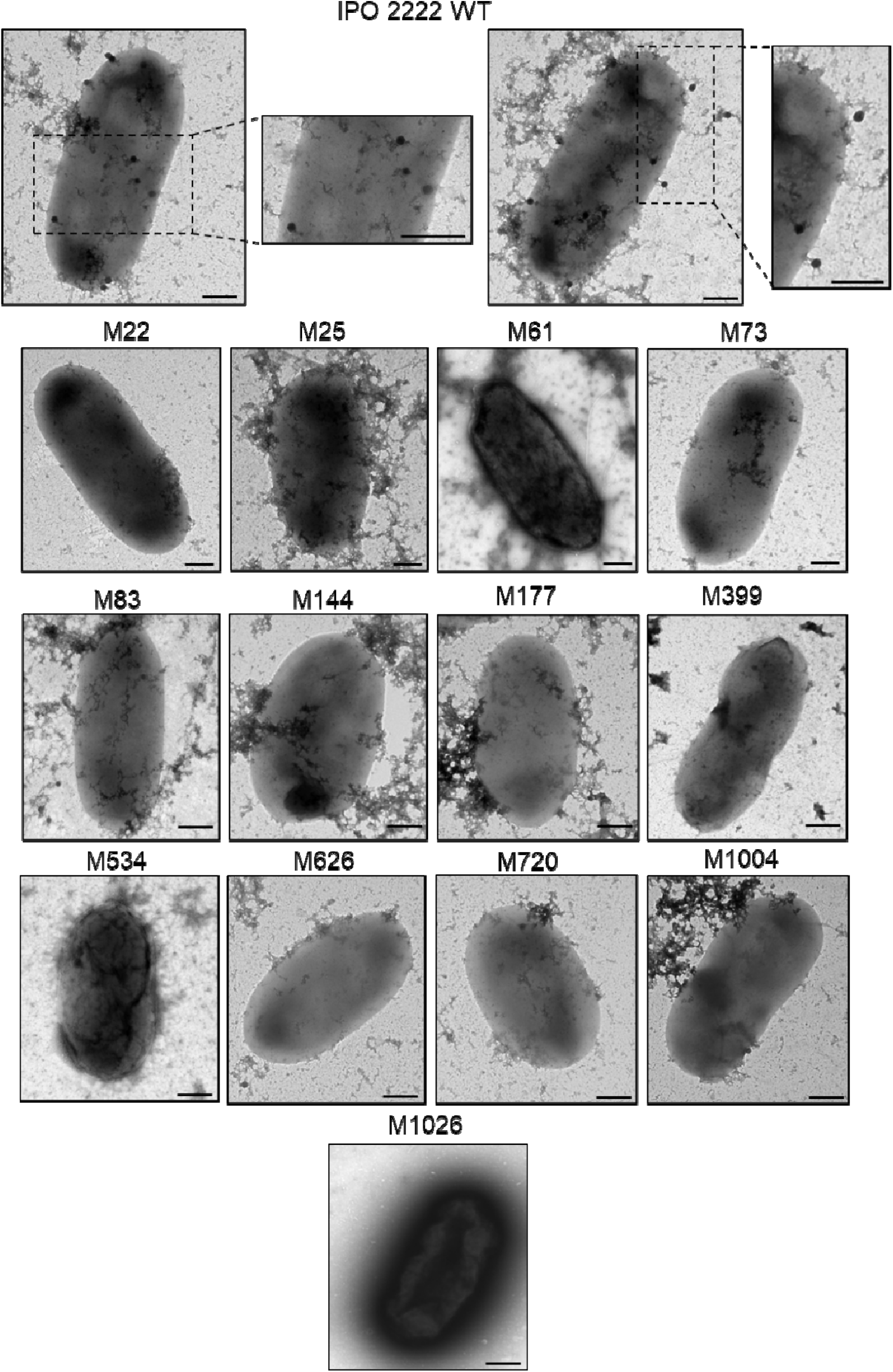
Visualization of adsorption of vB_Dsol_D5 (ΦD5) particles to *D. solani* IPO 2222 WT and 13 phage-resistant mutants by transmission electron microscopy (TEM). Bacterial cells and phage particles were mixed at MOI of 10 and incubated for 20 min at room temperature (ca. 20-22 °C) to allow the phages to attach to host bacterial cells. At least 10 individual images were gathered for each analyzed strain, and the experiment was repeated once (two biological replicates of the assay). Representative photos are shown. Scale bar – 200 nm.

**Figure 4.**
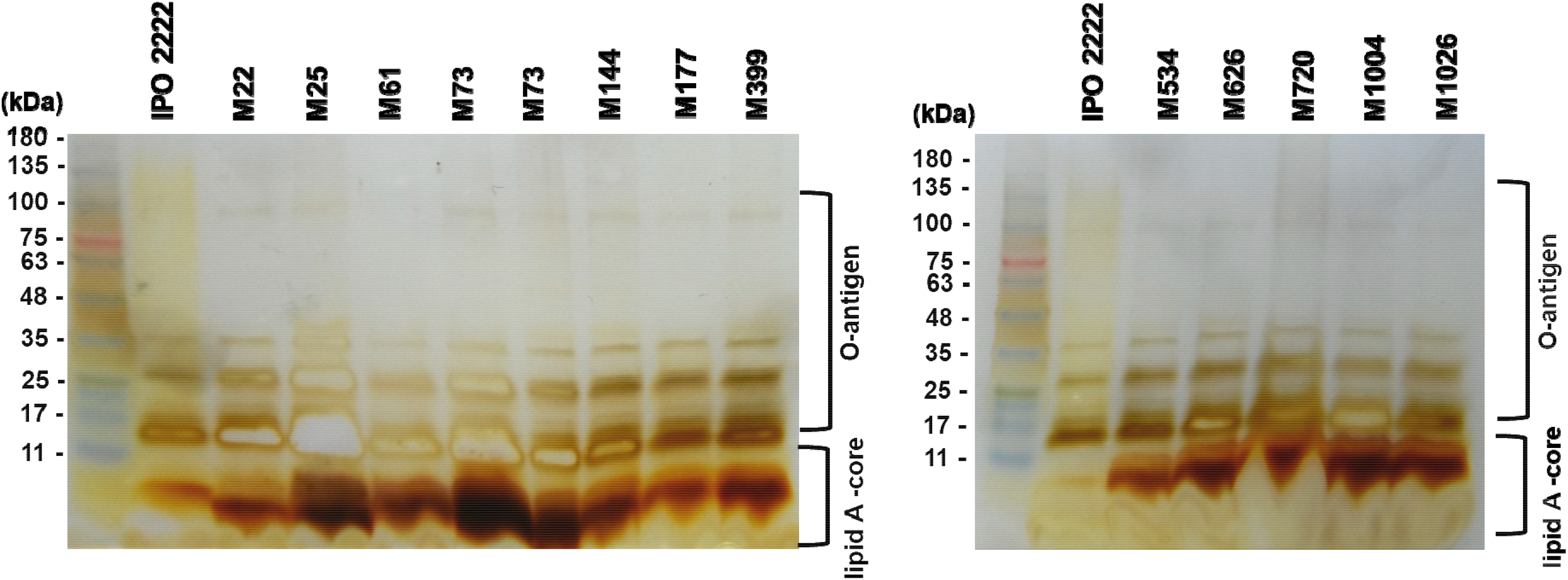
Characterization of lipopolysaccharide (LPS) isolated from *D. solani* IPO 2222 WT and 13 phage-resistant Tn5 D. solan animutants. SDS-PAGE was made using a gradient (4-20%) polyacrylamide gel (Biorad), and the LPS components were visualized u using silver staining. The size marker (11-245 kDa, Perfect Tricolor Protein Ladder, EURx, Poland) is shown in the first lane of both gels els.

### Mass spectrometry analysis of *D. solani* IPO 2222 and ΦD5-resistant Tn5 mutants’ surface proteins

The intact cells MALDI technique was used to analyze the profile of the membrane proteins present on the surface of phage-resistant mutants. Viable cells for each Tn5 mutant and IPO 2222 WT were applied directly from the fresh TSA plate to the MS measurement plate and covered with an appropriately prepared matrix mixture as described above. Recorded MS spectra were analyzed in groups corresponding to Tn5-mutated loci found in phage-resistant mutants (Tab. 1, Fig. 1). The spectra in the range of 5 000 to 20 000 m/z are shown in Supplementary Figure 1, and their approximations for the range of 5 000 to 9 400 m/z are in Supplementary Figure 2.

All the samples showed significant similarity to *D. solani* IPO 2222 WT strain. For all mutants, except for M61, which presents a minor signal spectrum, the pattern in the m / z from the 6000 to 6600 range was very well preserved. The most intense signals at 9534.6 m/z and two distinct ones at 6522.1 and 7203.2 m / z are visible in each of them as well. One can see changes between the intensity of the last two mentioned signals, but there is no regularity in the analyzed groups (one of them prevails in the group in each mutant). However, subtle differences can be found between individual groups or single variants versus wild-type cells. For example, the M22 and M25 variants have clearly distinguished two signals at 7266.9 and 7287.2 m / z, which have a much lower intensity (Supplementary Figure 1). Similarly, intense signals are distinguished for variants M177, M83 and M626. In all cases, proteins corresponding to signals above 10,000 m/z are hardly visible. Most of them are broad, low-intensity signals. Most variants present fewer peaks in this range than the wild variant. The signal at 10665.2 is present in every MS spectrum. The preceding 10,300 m / z signal seen by IPO2222 disappears in the remaining cells. It is visible only for variants from the M83/399/534 group. They also give relatively distinct remaining higher masses between 15,000 to 17,000. Of the remaining variants, the peak at 16,222.7 is marked for M61 and M177. In the range of about 12,000 m/z, two signals, 12 290.2 and 12345.0, are visible for the WT. In the case of the others, the first signal is visible mainly.

### Analyses of the cell aggregation

In repeatable experiments, an average IPO 2222 WT (control) sedimentation (due to cell aggregation) of ca. 7% was observed after 24 h incubation. In repeatable experiments of the 13 phage-resistant mutants analyzed for cell aggregation, only two mutants (M22 – sedimentation of ca. 10% cells and M25 – sedimentation of ca. 10% cells) did not express significantly accelerated sedimentation due to elevated cell aggregation in comparison with the IPO 2222 WT strain (Figure 6). All other 11 mutants expressed statistically significant increased sedimentation under experimental conditions; the highest level of sedimentation was observed in the case of mutants M83, M177, M399, M534 and M1004, in which on average between 81 and 92 % of cells sedimented to the bottom of the culture flasks after 24 h incubation (Figure 6).

### Analyses of the cell surface of phage-resistant Tn5 *D. solani* mutants using scanning electron microscopy (SEM) and atomic force microscopy (AFM)

In SEM analyses, IPO2222 WT cells were characterized by a gently folded cell wall and rounded cell poles (Fig. 5). Contrary, all phage-resistant mutants analyzed in this study had a more rough cell wall surface than IPO2222 WT (Fig. 5). The degree of the cell wall folding varied per phage-resistant mutants tested. A clearly spongy surface characterized M22 and M25 cells, whereas mutants M61, M144, M534 and M1004 had visible longitudinal furrows on the cell surface (Fig. 5). Mutants M73, M83, M177 and M626 were characterized by a lobular shape of the cell surface. The pointed cell poles and longitudinal surface folding were specific to the M399 mutant. M720 cells, apart from the longitudinal furrows in the wall, were characterized by the presence of forms bent even at an angle of 90 °. The M1026 cells had clearly shorter forms than the others and developed compact aggregates (Fig. 5).

**Figure 5.**
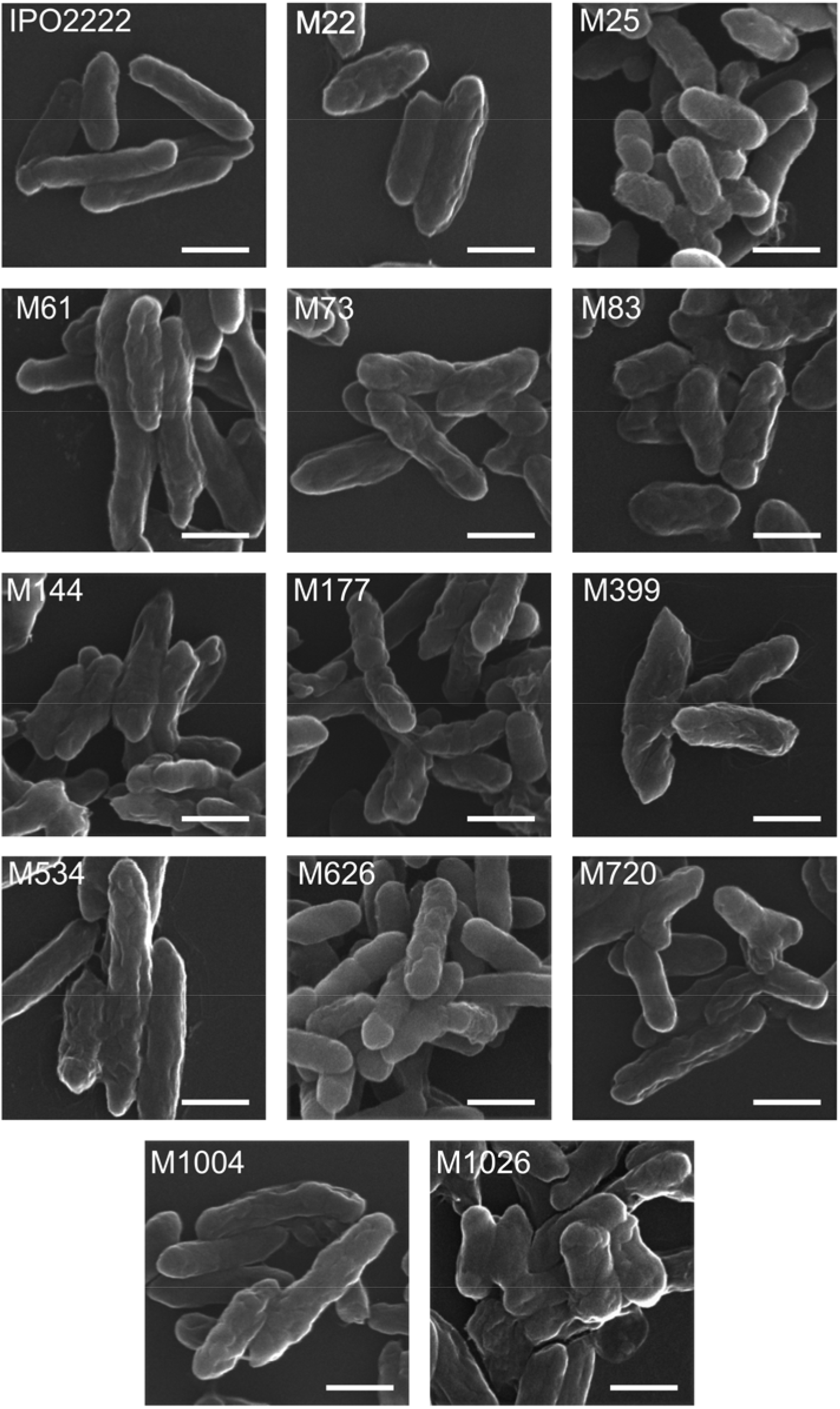
SEM imaging of D. solani IPO 2222 WT and 13 phage-resistant mutants. Scale bar corresponds to 1 µm.

Based on the SEM results, we have chosen two mutants (M399 and M1004) for AFM analyses. The chosen mutants both possessed a more rough surface area than all other tested strains, as evidenced in SEM analyses (Fig. 5). Mutant M399 was characterized by a less regular surface than the IPO 2222 WT strain. The second selected mutant, M1004, had a folded surface significantly greater than both IPO 2222 WT and M399 mutant. Due to the differentiation of the cell surface, the profile of two sites of each cell was analyzed. AFM imaging of the *D. solani* IPO2222 WT strain showed a regular and symmetrical cell profile (Fig. 6). Slight folds characterized cell wall surface both at three-dimensional image and height profiles, as shown in (Fig.6, panels A1-A3). The mutant M399 and M1004 cells’ topography was visibly different from that of the *D. solani* IPO2222 WT cells. Distinct folding, irregular cell wall surface and furrows were visible in the height profiles of the selected mutant cell (Fig. 6, panels B1-B3, C1-C3). Moreover, the surface roughness of the *D. solani* IPO2222 WT was lower (average roughness, Ra=3.05 nm) than the average roughness of the surfaces of *D. solani* phage-resistant mutants M399 (Ra =4.41 nm) and M1004 (Ra=4.89).

**Figure 6.**
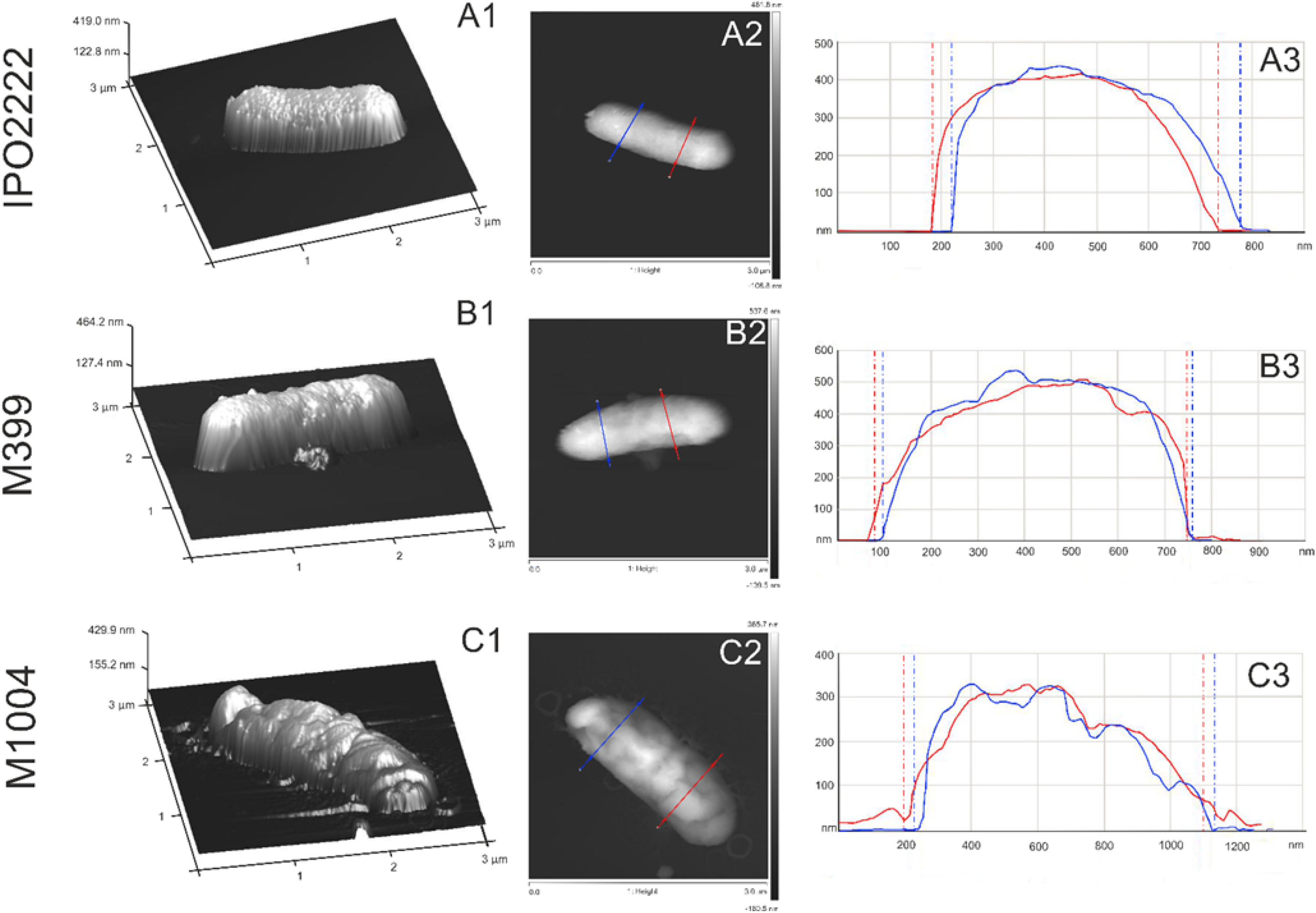
Atomic force microscopy (AFM) imaging of D. solani IPO 2222 WT and selected phage-resistant mutants M399 and M1004. (panel A1–A3) – cell-surface *D. solani* IPO2222 WT; (panel B1–B3) – cell surface of phage-resistant mutant M399; (panel C1–C3) – cell surface of phage-resistant mutant M1004. (panel A1, B1, C1) –three-dimensional images of *D. solani* WT and mutant cells; (panel A2, B2, C2) –height mode images of cells; (panel A3, B3, C3) – height profiles of cells, the profiles were made along the lines shown in the height images.

### Phenotypes of *D. solani* phage-resistant mutants

The phage-resistant Tn5 mutants were tested for phenotypes that may be important for the ecological fitness and virulence of *D. solani* under natural and agricultural settings. No differences were found among 13 phage-resistant mutants and WT strain in the most metabolic phenotypes examined using BIOLOG GEN III and Ecoplate phenotypic microarrays. The ΦD5-resistant mutants differed from the WT strain in a total of only 3 features out of 94 testes with GEN III plates and from 31 tested using Ecoplate microarrays. Mutants M73, M144, M626 and M720 (locus: O-antigen export system (LPS transport system) ATP-binding protein) lost the ability to utilize D-cellobiose as a sole carbon source, and mutants M22 and M25 (locus: glycosyltransferase family 1) became susceptible to troleandomycin. In addition, mutants M73, M83, M144, M177, M399, M534 M626, M720 and M1004 (loci: *wzt* (in M73, M144, M626 and M720), putative *wbeA* (in M83, M399 and M534) and *fcl* (in M177 and M1004)) gained the ability to utilize inosine, a feature absent both in IPO 2222 WT and 4 other phage-resistant mutants (M22, M25, M61 and M1026) tested.

Likewise to the IPO 2222 WT (which produced cavities on CVP, produced proteases and degraded carboxymethylcellulose and polygalacturonic acid, it did not produce siderophores and was unable to grow on TSA medium supplemented with 5% NaCl), all phage resistant Tn5 mutants maintained these phenotypes. Furthermore, phage-resistant mutants analyzed in this study retained the ability to form biofilm. Except mutants M22 and M25, which produce a similar amount of biofilm as the WT strain, all other mutants produced biofilm with statistically higher abundance than the IPO 2222 WT. Using transmission electron microscopy (TEM), no considerable difference in cell morphology and diameter was noted in the phage-resistant mutants compared to the IPO 2222 WT strain. Likewise, all phage-resistant mutants exhibited similar colony morphology and colony diameter to that of the IPO 2222 WT strain. None of the tested phage-resistant mutants differed significantly in their average generation times in rich (TSB) medium. However, two mutants, M22 and M25, expressed statistically significant delayed growth in the minimal medium (M9+0.4% glucose) compared to the growth of IPO 2222 WT strain (Fig. 5). The analyses of mutants’ growth at six different temperatures (4, 8, 15, 28, 37 and 42 °C) did not indicate any growth differences compared to that of the IPO 2222 WT strain. All tested strains grew at 8, 15, 28 and 37 °C but were unable to grow at 4 and 42 °C. The growth of the mutants at acidic pH 5.0 and basic pH 10.0 was similar to that of the IPO 2222 WT strain (data not shown). The wild-type IPO 2222 expressed both swimming and swarming motility, whereas mutants M83, M177, M399, M534 and M1004 were nonmotile both when tested for swimming and swarming phenotypes. All remaining Tn5 mutants expressed significantly reduced swimming motility compared to the *D. solani* IPO 2222 WT strain. None of the 13 phage-resistant mutants displayed swarming motility in repeatable experiments. All mutants expressed similar susceptibility/resistance to all of the antibiotics tested as the IPO 2222 WT strain, except that all 13 Tn5 mutants expressed the resistance against kanamycin not observed in the wild type strain, which appeared as an effect of the transposon mutagenesis (=the neomycin resistance gene on the Tn5 transposon confers the cross-resistance to kanamycin). In repeatable experiments, all phage-resistant *D. solani* mutants were also significantly affected in producing extracellular polymeric substance (EPS). Five mutants (M22, M25, M83, M399 and M534) expressed reduced concentration of EPS in comparison with the WT strain (= ca. 8 to 23% of EPS produced by the wild type strain). In contrast, elevated concentration of EPS in comparison with IPO 2222 WT strain (= ca. 175-210% of the EPS produced by IPO 2222 WT) was recorded for 8 other mutants tested (M61, M73, M144, M177, M626, M720, 1004 and M1026).

### Survival of phage-resistant Tn5 mutants on potato leaves

In replicated experiments, survival of 13 ΦD5-resistant *D. solani* Tn5 mutants and *D. solani* strain IPO 2222 WT on the adaxial surface of detached potato (*S. tuberosum*) leaves was monitored for 14 days (Fig. 6). The numbers of bacteria differed widely per plant and treatment but were not statistically different between experiments 1 and 2, and thus the two experiments were analyzed together. In both experiments, at 0 dpi (at the stage of leave inoculation), the average bacterial populations of 10^5^ – 10^6^ CFU per leaf were found in all treatments. No additional statistically significant changes in the *D. solani* IPO 2222 WT population at 14 dpi compared to 0 dpi were observed in both experiments (Fig. 7). In contrast, both experiments recorded a statistically significant decrease of bacterial populations on leaves for all 13 phage-resistant D. solani mutants. During two weeks (at 14 dpi), *D. solani* Tn5 mutants populations dropped on average from ca. 10^5^ – 10^6^ CFU to ca. 10^2^ – 10^3^ CFU per leaf (Figure 6). In both experiments, no pectinolytic, cavity-forming bacteria were found on the adaxial side of non-inoculated leaves (negative control).

**Figure 7.**
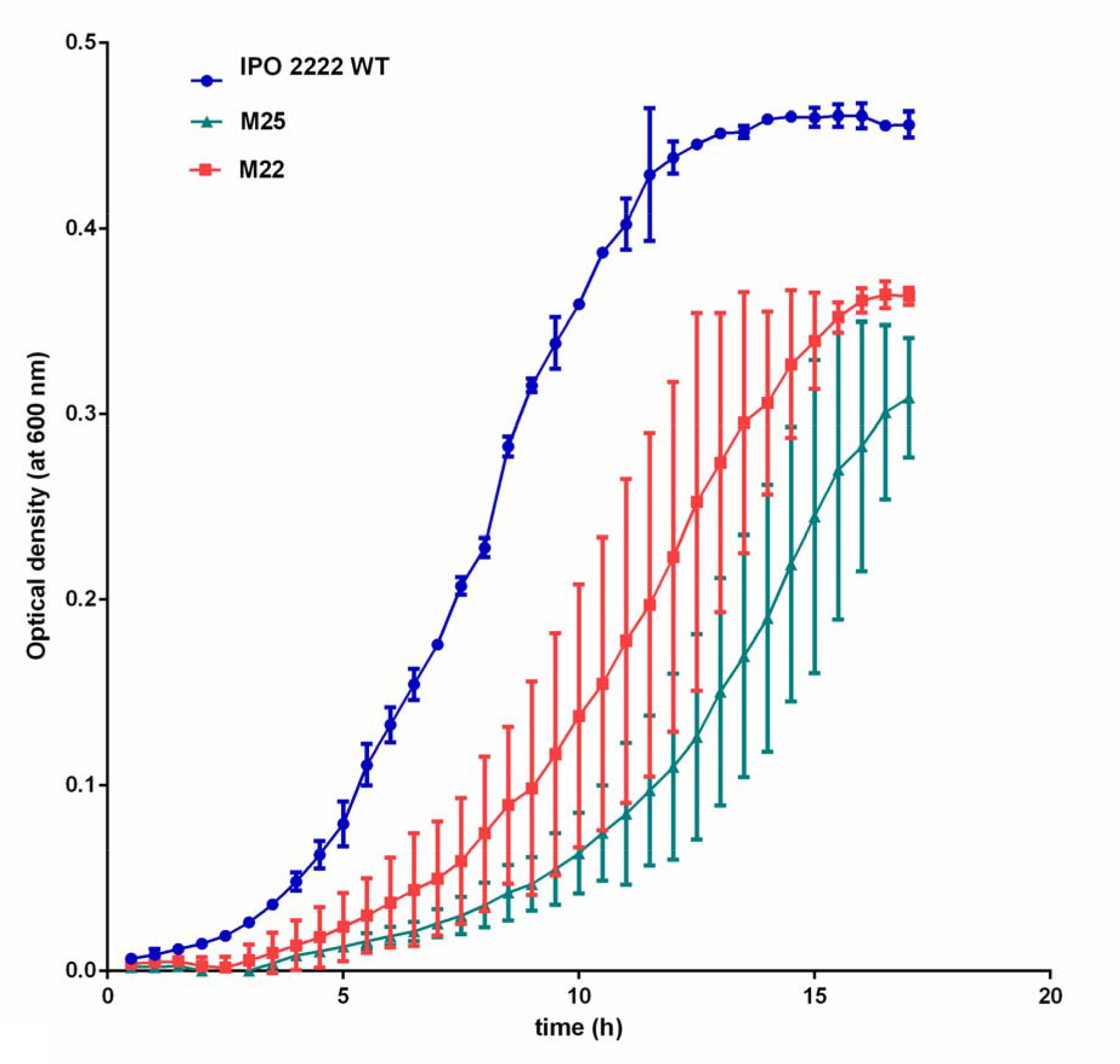
Growth of *D. solani* IPO 2222 WT and phage-resistant Tn5 *D. solani* mutants M22 and M25 in minimal growth medium (M9+0.4% glucose). The experiment was performed in two biological replicates containing two technical replicates each (n=4). The results were averaged for presentation. The bars show standard deviation (SD).

### Virulence of ΦD5-resistant *D. solani* Tn5 mutants in potato tubers *in vitro* and potato plants grown under growth chamber conditions

Although all 13 phage-resistant mutants expressed some level of virulence when applied to the interior of potato tubers by stab-inoculation, they all were significantly affected in their ability to macerate potato tuber tissue compared with IPO 2222 WT strain (Fig. 7).

Likewise, all 13 *D. solani* Tn5 mutants were significantly less virulent when tested for the development of blackleg symptoms on potato plants grown in potting compost infested with bacteria. In two separate phytochamber experiments, most potato plants inoculated with phage-resistant Tn5 mutants did not express any blackleg symptoms. In contrast, between 80 and 100% of the plants inoculated with IPO 2222 WT strain (positive control) developed severe and typical blackleg symptoms, which led even to the death of several infected plants. As expected, no symptoms were observed at any time point in plants inoculated with sterile Ringer’s buffer (negative control) in both growth chamber experiments.

The population size of the IPO 2222 WT differed significantly between positive control plants. However, the viable bacterial cells were recovered from all (100%) inoculated plants in both experiments at 14 days post-inoculation at densities ranging from 1000 to 15000 CFU g^-1^ of the stem tissue. The stems of negative control plants did not harbour *D. solani* cells. A much lower incidence of recovery of phage-resistant mutants from stem after soil infestation was observed in both experiments. Phage-resistant strains were incidentally detected in only 4 plants expressing typical blackleg symptoms; of ten plants per treatment, one plant harboured M73 mutant (ca. 1000 CFU g^-1^ of stem tissue), one M144 mutant (ca. 1000 CFU g^-1^), one M626 mutant (ca. 1100 CFU g^-1^), and one M720 mutant (ca. 1000 CFU g^-1^). None of the other tested mutants was detected inside stems of inoculated plants in both experiments (Fig. 8).

**Figure 8.**
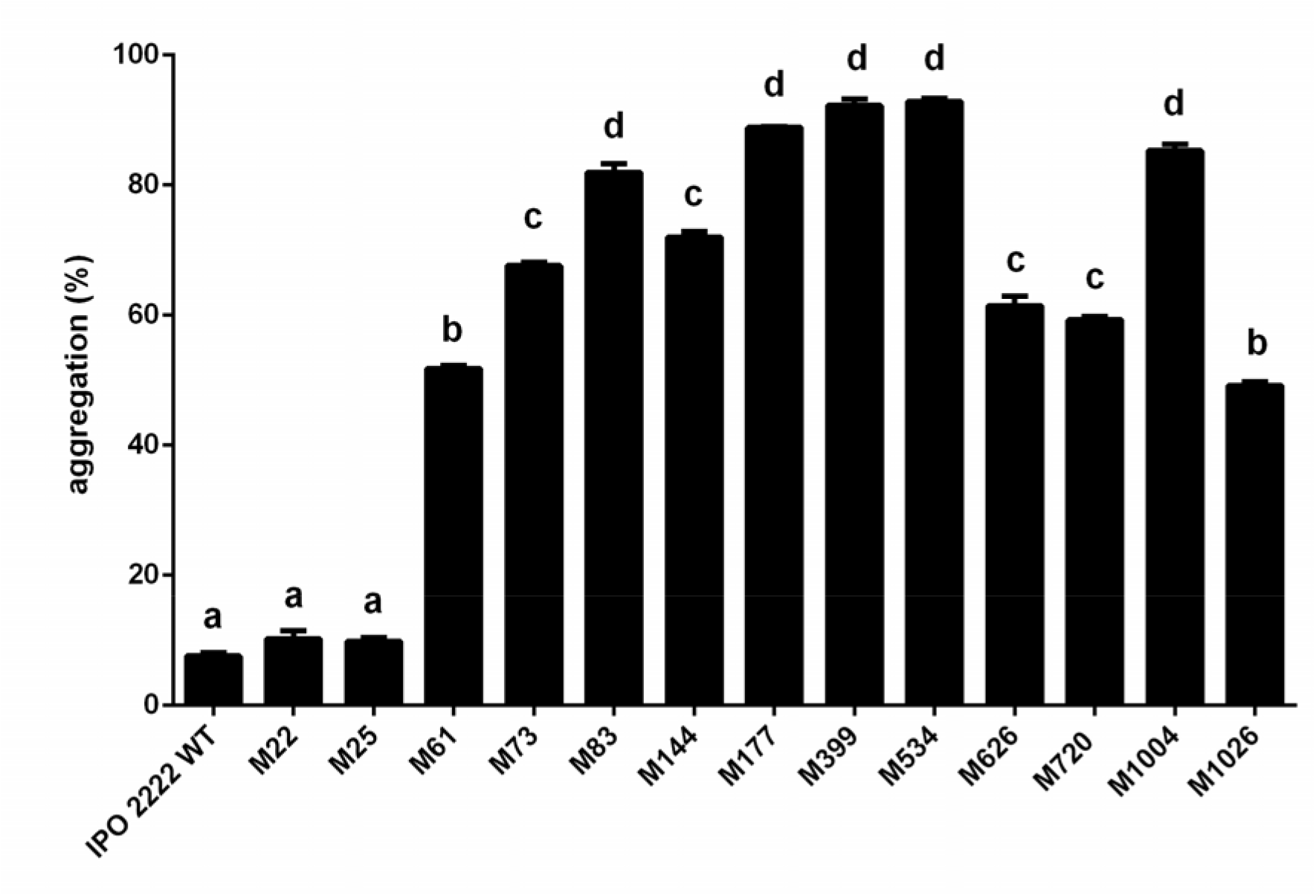
Percentage of aggregation of cells of phage-resistant Tn5 *D. solani* mutants. The percentage of aggregation was quantified from the change in optical density (OD600) over 24 h. Percentage aggregation (sedimentation) was measured as follow: %A = 1 (OD60024h/OD600 0h), where: %A – percentage of aggregation (sedimentation), OD6000h – OD of bacterial culture at time 0 h, OD60024h – OD of bacterial culture at time 24 h

### Construction of *D. solani* strains tagged with GFP or DsRed for competition experiments

Transformation of *D. solani* IPO 2222 WT and Tn5 phage-resistant mutant strains with pPROBE-AT--*gfp* and/or pRZ*-*T3*-dsred* plasmids resulted in ca. 20 to 50 transformants per strain obtained per transformation event. One bacterial colony expressing a high fluorescent signal was collected and used for further analysis in each case. At least three repeated transfers of individual transformants on TSA supplemented with antibiotic (ampicillin or tetracycline) were done to confer the stable expression of GFP or DsRed, respectively, in each transformed strain (Supplementary Table 1.). Fluorescently-labelled bacterial variants displayed similar growth characteristics in liquid media to their parental strains (data not shown), indicating that the growth of the bacterial strains was not altered either by the presence of plasmids carrying *gfp* or *dsred* genes or by expression of fluorescent proteins in transformants (data not shown).

### Competition of marker *D. solani* WT and Tn5 mutant strains tagged with GFP or DsRed on potato tubers

In two separate phytochamber experiments, the competition of fluorescently-labelled IPO 2222 WT and phage-resistant Tn5 mutants was assessed after 72 h incubation of stab-inoculated tubers and under conditions provocative for development of disease symptoms (28 °C, ca. 80-90% relative humidity (RH)). The numbers of bacteria differed broadly per tuber and per treatment applied but were globally not statistically different between experiments 1 and 2, and consequently, the results from both experiments were analyzed together. After 72 h incubation, the average populations of *D. solani* IPO 2222 WT in singly inoculated tubers (control) reached between 10^6^ and 10^7^ CFU g^-1^ of tuber tissue. The average populations of the WT strain in tubers jointly inoculated with phage-resistant mutants were slightly reduced compared to the *D. solani* populations in positive control tubers and were ca. 5 x 10^5^ CFU g^-1^ of tuber tissue. In potato tubers inoculated with different *D. solani* Tn5 mutants (mutant control) alone, the bacterial populations of all mutants were significantly reduced compared to the populations of IPO 2222 WT but fairly similar to each other and were ca. 10^3^ – 10^4^ CFU g^-1^ of tuber tissue depending on the analyzed Tn5 mutant (Fig. 9). The average populations of the Tn5 mutants in tubers inoculated jointly with IPO 2222 WT (treatments) decreased compared to the treatments containing Tn5 mutants alone. They reached only ca. 10 – 5 x 10^2^ CFU g^-1^ after 72 h incubation (Fig. 9).

**Figure 9.**
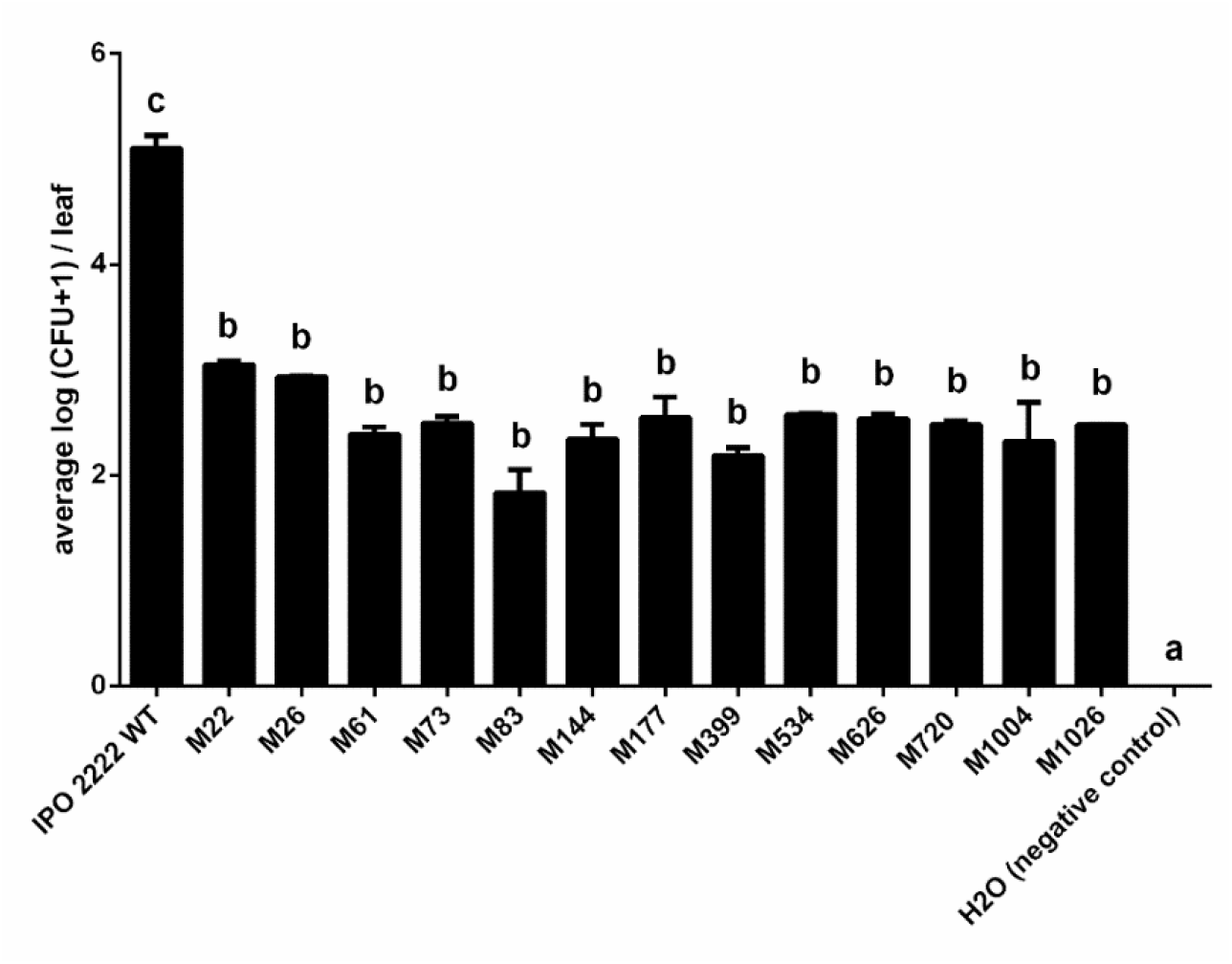
Populations of *D. solani* IPO 2222 WT and 13 phage-resistant Tn5 *D. solani* mutants on the adaxial surface of *S. tuberosum* detached leaves at 14 days post-inoculation. In replicated experiments, 20 leaves (10 leaves per experiment per mutant) were samples at two time points (0 [control] and 14 days post inoculation). Both at 0 and 14 dpi, five leaves (third to six from the shoot terminal) spray-inoculated with 2 ml of 10^6^ CFU mL^-1^ of bacterial suspension (WT or individual Tn5 mutant) in 1/4 Ringer’s buffer and incubated adaxial side up on 0.5% water agar in square plastic Petri dishes (100 x 100 mm) were sampled. At 0 and 14 dpi, five randomly chosen leaves from five randomly chosen Petri dishes were collected and individually shaken in 10 ml Ringer’s buffer in 50-mL Falcon tubes at 50 rpm at room temperature for 30 min to wash bacterial cells off the leaf surface. The serial diluted leaf washings were plated in duplicates on CVP supplemented with 200 µg mL^-1^ cycloheximide (for isolation of IPO 2222 WT) or on CVP containing 50 µg mL^-1^ neomycin ad 200 µg mL^-1^ cycloheximide (for isolation of phage-resistant Tn5 mutants). Inoculated plates were incubated at 28 °C for 24-48 h, and the resulting colonies were counted. The results were averaged for presentation. The bars show standard deviation (SD).

**Figure 10.**
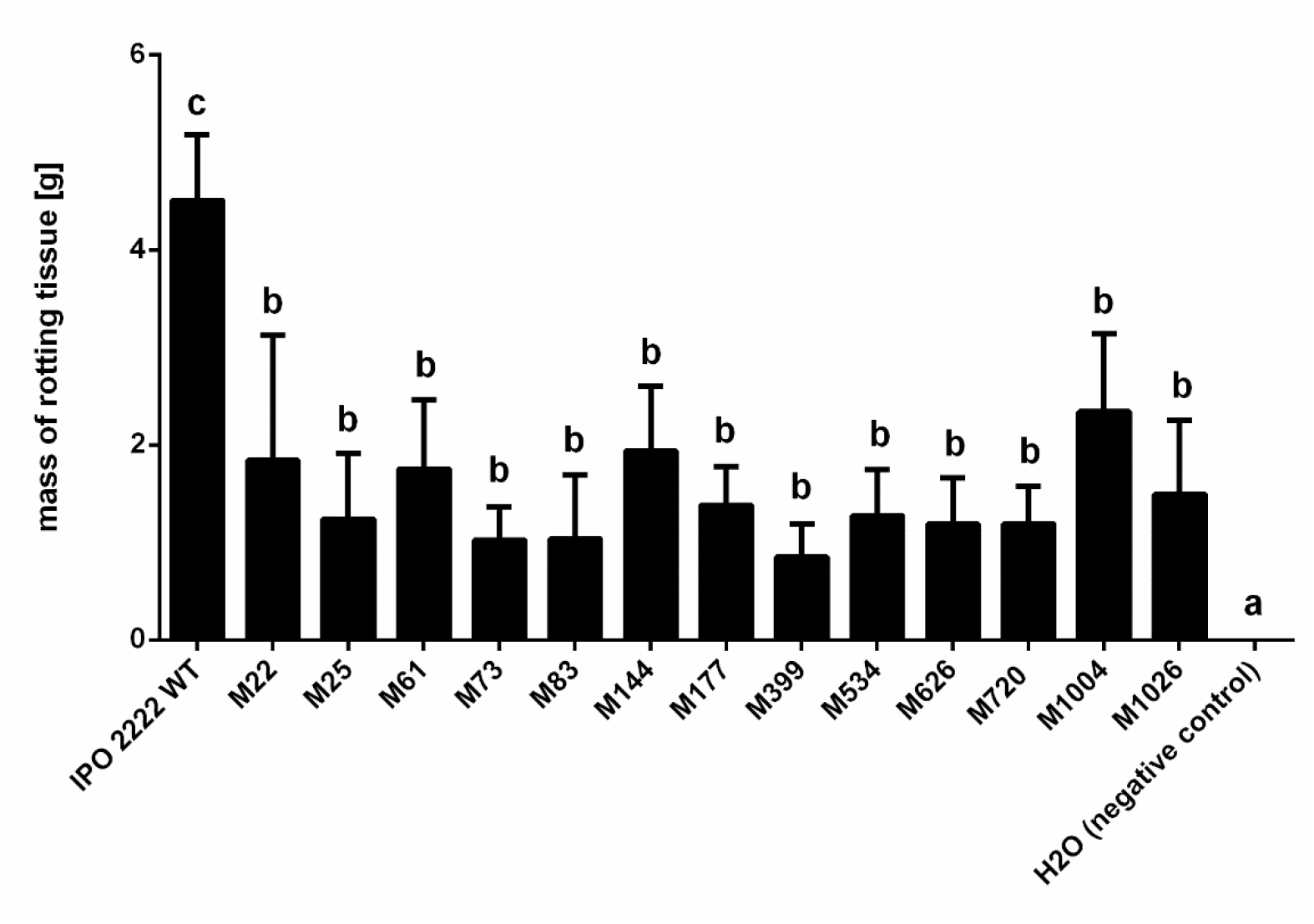
Maceration of potato tuber tissue by *D. solani* IPO 2222 WT and 13 phage-resistant. Tn5 *D. solani* mutants. Five individual potato tubers were inoculated with a given phage-resistant bacterial mutant using a whole tuber injection method (stab inoculation) (Czajkowski et al., 2012b; Czajkowski et al., 2017b). Bacterial strains were grown in TSB (WT strain), or TSB supplemented with 50 µg mL^-1^ of neomycin (Tn5 mutants) for 24 h at 28 °C. After incubation, bacterial cultures were separately collected, washed two times with 1/4 Ringer’s buffer and resuspended in the initial volume of Ringer’s buffer. Optical density (OD600=0.1) was used to normalize the number of bacterial cells in all treatments (ca. 10^8^ CFU mL^-1^). Surface-sterilized potato tubers were stab-inoculated with sterile yellow pipette tip filled with 50 µl of bacterial suspension (treatments) or sterile Ringer’s buffer (negative control). IPO 2222 WT was used as a positive control. Vertical lines represent standard deviation (SD).

**Figure 11.**
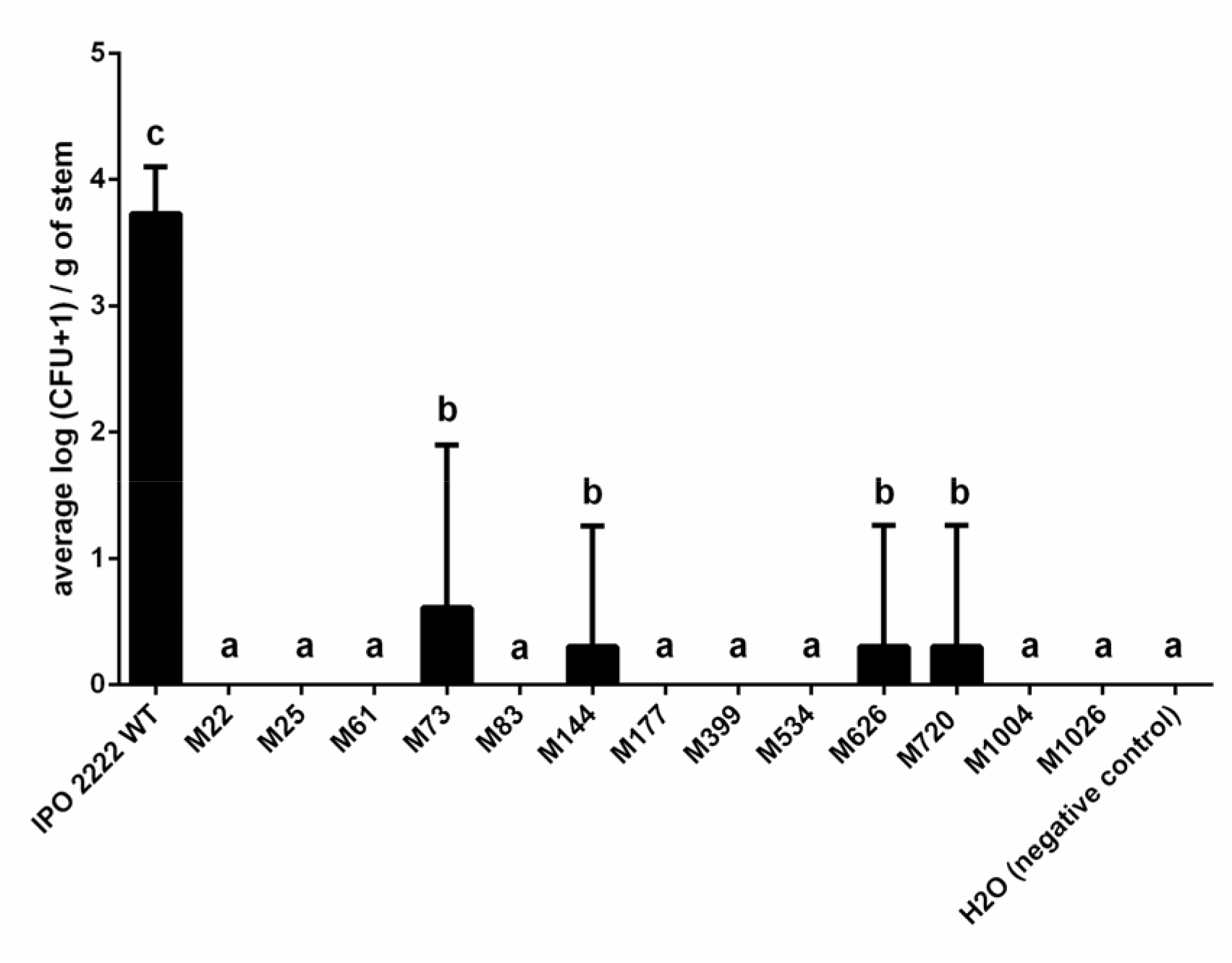
Population size of D. solani IPO 2222 WT and 13 phage-resistant Tn5 D. solani mutants inside stems of potato plants after inoculation into the soil. Results were considered significant at *p*=0.05, and the pair-wise differences were obtained using the t-test. The means that do not share the same letters above each bar differ. Vertical lines represent standard deviation (SD).

**Figure 12.**
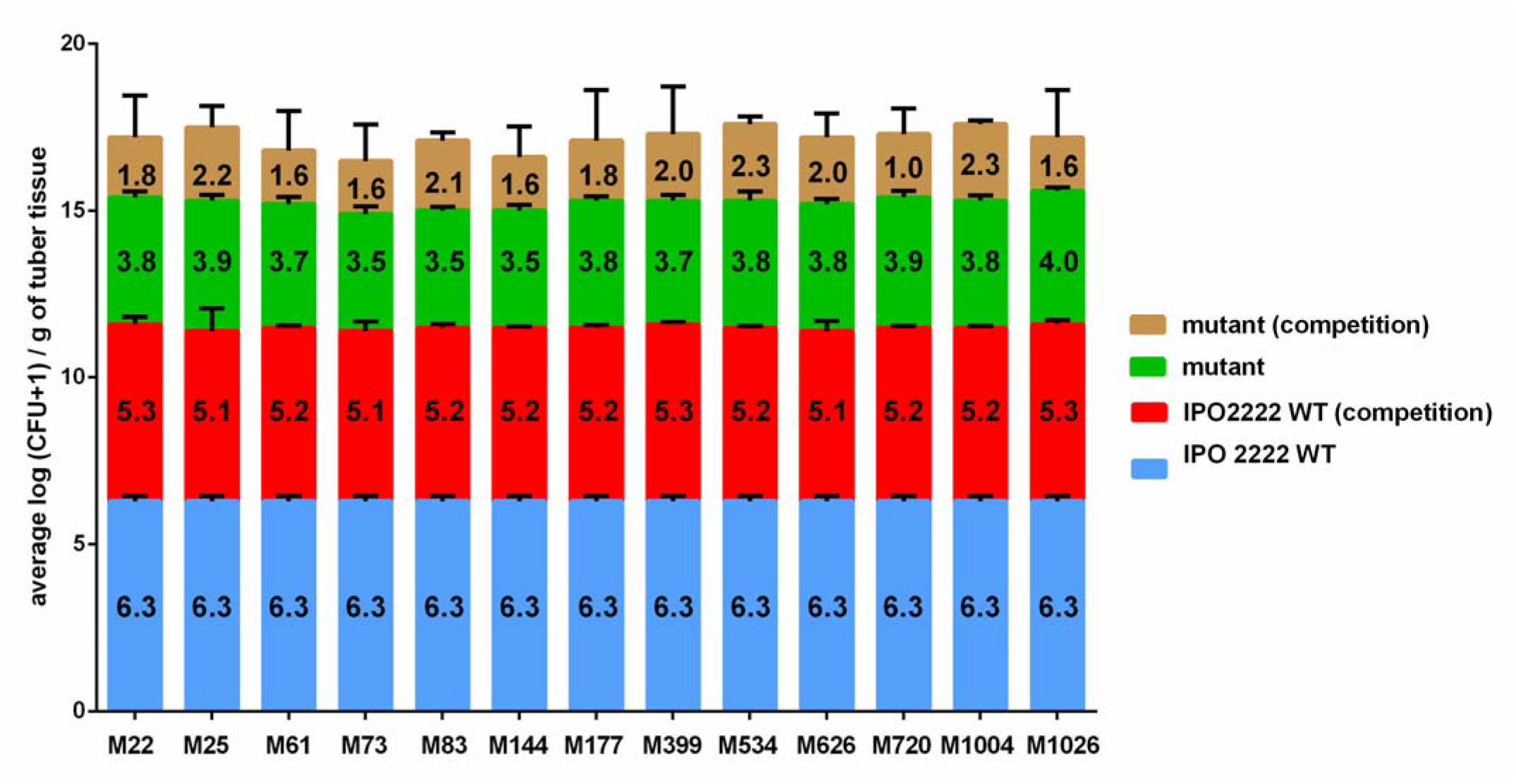
Competition of *D. solani* IPO 2222 WT and phage-resistant *D. solani* Tn5 mutants on potato tubers. Potato tubers were inoculated either with fluorescently labelled *D. solani* IPO 2222 WT, fluorescently labelled individual phage-resistant *D. solani* Tn5 mutants or co-inoculated with WT strain and individual Tn5 mutant. In the first experiment, a GFP-tagged *D. solani* IPO 2222 WT and DsRed-tagged phage resistant *D. solani* mutants were used. In contrast, DsRed-tagged IPO 2222 WT and GFP-tagged phage resistant D. solani mutants were applied in the second experiment. Inoculated tubers were kept under conditions that promote rotting (28 ° and 80-90% relative humidity). Three days (72 h) post inoculation tubers were sampled and analyzed for the presence of fluorescently tagged bacteria using pour plating. The GFP and DsRed positive colonies were counted. The experiment was replicated one time with the same setup, and the results from both repetitions were averaged for analyses. The numbers inside bars represent the average log (CFU+1/g of tuber tissue) for the respective treatment. Vertical lines represent standard deviation (SD).

## Discussion

This study is, to our knowledge, one of the first to examine the link between phage resistance and the fitness of the plant pathogenic *D. solani* strain IPO 2222 in its natural environment (*S. tuberosum* plants and tubers). Our main finding is that the cost of resistance of *D. solani* strain IPO 2222 to lytic phage ΦD5 depends on the environmental context. Although IPO 2222 resistance to ΦD5 did not affect most of the phenotypes of *D. solani* phage-resistant Tn5 mutants screened *in vitro*, all tested mutants were significantly impacted in their ability (i) to survive on the plant surface and (ii) cause disease symptoms in potato plants and tubers. Unfortunately, to date, the mechanisms governing the interaction of SRP bacteria and their lytic bacteriophages, as well as the ecological context of such associations, remain insufficiently explored (Czajkowski, 2016; Toth et al., 2021). This study used a Tn5-based random transposon mutagenesis method to identify *D. solani* mutants resistant to infection caused by lytic phage ΦD5. In addition, phage-resistant *D. solani* mutants were assessed for phenotypes likely to be involved in the ecological success of the pathogen in the plant environment. This approach allowed us to investigate the tradeoff hypothesis in which the ΦD5 resistance cause fitness disadvantages for the phage-resistant *D. solani* IPO 2222 variants during infection.

Screening the collection of 1000 *D. solani* Tn5 mutants allowed us to find 13 different bacterial mutants resistant to phage ΦD5. Surprisingly, all 13 Tn5 insertions were located multiply times in the small pool of *D. solani* genes (=5 individual loci, from 2 to 4 independent Tn5 insertions per mutant per particular locus of interest). Furthermore, we found that the mutated genes are the parts of only two operons (Fig. 1, Table 1), both previously described as involved in the envelope status of Gram-negative bacteria, including bacteria belonging to *Enterobacteriaceae* and *Pectobacteriaceae* families (Holt et al., 2020). Specifically, these operons are participating in the biosynthesis of the core region of LPS together with the biosynthesis of O-antigen of LPS in Gram-negative bacteria (Fig. 1) (Schnaitman and Klena, 1993; Ranjan et al., 2021). As no phage resistant mutants analyzed in this study had Tn5 insertions in other *D. solani* genes and operons, we believe bacterial envelope components (*viz*. LPS, EPS and extracellular capsule (CPS)) encoded by the operons mentioned above are essential for ΦD5 interaction with IPO 2222 cells. To further acquire more information about the LPS in the context of phage resistance, we compared the lipopolysaccharides from phage-resistant mutants with that of the IPO 2222 WT strain. Surprisingly, the characterization of LPSs by SDS-PAGE did not reveal apparent structural differences between the LPS of the WT strain and Tn5 mutants that could immediately explain the ΦD5-resistant phenotype.

Modification of bacterial envelope is one of the most common strategies of phage evasion in Gram-negative bacteria (Bohannan and Lenski, 2000; Mangalea and Duerkop, 2020). To date, however, only limited data exist on how frequently such a mechanism is employed by SRP and especially by *D. solani* to prevent phage infection. In this study, 11 of 13 phage-resistant mutants expressed a complete inhibition, whereas the other two (M177 and M1004) substantially reduced phage adsorption. This observation straightforwardly connects the phage adsorption and the status of the *D. solani* envelope. In our former study, in which we analyzed the interaction of *P. parmentieri* SCC 3193 with its lytic bacteriophage ΦA38, we found out that the ΦA38 requires intact LPS to infect its host and that the alterations in LPS synthesis protected *P. parmentieri* SCC 3193 from viral infection (Bartnik et al., 2021). Similar observations linking phage-resistance and cell surface status were done for other SPR bacteria, including *P. atrosepticum* strain SCRI 1043, *P. carotovorum* strain Pcc27 and *P. brasiliense* strain F152 (PB29) (Evans et al., 2010; Kim et al., 2019; Lukianova et al., 2019) but not for *D. solani* strain IPO 2222.

Knowledge of the ecological functions of bacterial envelopes, including LPS, EPS and CPS polysaccharides (capsule), comes in majority from studies done on human and animal pathogenic bacteria (Costerton et al., 1981; Beveridge and Graham, 1991). In contrast, the role of bacterial surface status in the interaction of plant pathogens and their plant hosts is comparatively much rarer assessed (D’Haeze and Holsters, 2004). In mutants M22 and M25, the gene coding for glycosyltransferase linked to extracellular polysaccharide synthesis was disrupted by the presence of the Tn5 transposon. Glycosyltransferases are involved in LPS and EPS synthesis in plant pathogenic and plant beneficial bacteria (Santaella et al., 2008; Li and Wang, 2012). In-plant pathogenic *Xanthomonas citri* subsp. *citri* (Xac) glycosyltransferase was reported to be required for EPS and LPS production. Glycosyltransferase defective Xac mutants were impaired in biofilm and EPS production, demonstrated delayed growth and expressed increased susceptibility to environmental stresses than the wild type strain (Li and Wang, 2012). In plant-beneficial *Rhizobium* sp. YAS34, polysaccharide-associated glycosyltransferase, was involved in the synthesis of EPS, and it was critical for colonization of plant roots as their activity stabilized cell adherence during plant colonization (Santaella et al., 2008). These phenotypes were in the majority confirmed in this study in M22 and M25 mutants, which expressed reduced growth in minimal medium *in vitro*, reduced synthesis of EPS and decreased swimming and lack of swarming motility. The *cpsB* (*manC*) gene disrupted in mutants M61 and M1026 encodes mannose-1-phosphate guanylyltransferase /mannose-6-phosphate isomerase. *cpsB* was reported to be required for virulence of *Streptococcus penumoniae* and *Klebsiella pneumoniae* in mice (Morona et al., 2004; Lawlor et al., 2006). In-plant pathogenic *Erwinia amylovora*, a homolog of the *cpsB* gene, is a part of the GDP-fucose synthesis pathway involved in the synthesis of a capsular polysaccharide colanic acid.

Consequently, *E. amylovora* mutants lacking functional *cpsB* expressed avirulent phenotype (Geider et al., 1991). Furthermore, colanic acid was required for full virulence of soft rot *Pectobacterium* spp. close related to *Dickeya* spp. (Mohamed et al., 2015). In *Xanthomonas campestris* pv. *campestris* (Xcc), a homolog of *cpsB,* gene *xanA* is required for the synthesis of extracellular polysaccharide xanthan, involved in stress tolerance and required by Xcc both for attachment to plant surfaces and biofilm formation (Katzen et al., 1998). Here, *D. solani* phage-resistant mutants M61 and M1026 with disrupted locus coding for mannose-1-phosphate guanylyltransferase /mannose-6-phosphate isomerase expressed elevated biofilm formation, reduced swimming motility and increased EPS production. The *wzt* (*rbfB*) gene was disrupted in *D. solani* mutants M73, M144, M626 and M720. This gene encodes the O-antigen export system (LPS export system) ATP binding protein involved in the translocation of O-antigen across the inner cell membrane to the periplasm, attaching it to a lipid A core (Whitfield et al., 2020). In *Rhizobium tropici* CIAT899, *wzt* deletion mutant was affected in growth rate, motility, and it colonized plant roots to a lower frequency than the wild type strain (Ormeno-Orrillo et al., 2008). In *D. dadantii* A1828, *wzt* negative mutants express reduced survival under NaCl osmotic stress (Touze et al., 2004). In gliding bacterium, *Myxococcus xanthus* mutation in O-antigen export system (LPS export system) ATP binding protein leads to defects in social motility and colony formation under laboratory conditions (Bowden and Kaplan, 1998). In our study, all 4 mutants with Tn5 located in the *wzt* gene showed a similar phenotype but divergent from the phenotype of the WT strain: they express elevated EPS and biofilm formation and reduced ability to move. Gene *wbeA* encoding hypothetical protein (putative glycosyltransferase WbeA) was disrupted by Tn5 in the next three phage-resistant *D. solani* mutants (M83, M399 and M534) obtained in this study. The role of WbeA has not been assessed in detail in plant pathogenic bacteria yet. Likewise, the knowledge of the function of WbeA in the ecology and pathogenicity of SRP bacteria is limited (Ranjan et al., 2021). *wbeA* gene is a conserved member of the putative O-antigen LPS gene cluster present in members of the genus *Dickeya* spp. (Ranjan et al., 2021). Its glycosyltransferase activity and genomic location suggest that its involvement in LPS synthesis, together with another glycosyltransferase WbeB, present in the same operon (Ranjan et al., 2021). In our study, mutants M83, M399 and M534 expressed elevated biofilm formation, reduced EPS production and were nonmotile is swarming motility assay. In our study, the last two mutants, M177 and M1004 that were found to be ΦD5-resistant, had a mutation in gene *fcl* coding for GDP-L-fucose synthase. GDP-L-fucose synthase is required to synthesize extracellular polysaccharide colanic acid in *Escherichia coli* strain K-12 and other members of *Enterobacteriaceae* (Andrianopoulos et al., 1998). Fcl was reported to be involved in the synthesis of EPS in *Pectobacterium carotovorum* (Islam et al., 2019). The *fcl P. carotovorum* knockout mutant produced an elevated level of biofilm and expressed reduced virulence *in planta* (Islam et al., 2019). These phenotypes were similar to the phenotypes observed in *D. solani* phage-resistant mutants M177 and M1004.

To further explore the possibility that the surface status of *D. solani* cells is essential for ΦD5 infection, we assessed the morphology of IPO 2222 WT and the corresponding phage-resistant mutants using microscopic techniques. All phage-resistant mutants were indistinguishable from the WT strain in the colony and cell morphology. However, more detailed analyses done with SEM and AFM confirmed that all 13 phage-resistant mutants possess rougher and more irregular cell surfaces compared to the surface of the WT strain. In line with our observations, several other studies have already confirmed that bacterial mutants with rough envelope phenotype express resistance to viral infection (Qimron et al., 2006; Pagnout et al., 2019). Furthermore, envelope regularity and smoothness resulting from LPS, EPS and capsule cooperation are also among the key factors determining bacterial ability to form biofilms on different surfaces and survive in harsh environments (Montanaro and Arciola, 2000; Berne et al., 2018). Alterations of envelope properties may therefore increase antibiotic susceptibility, influence biofilm formation and reduce resistance to environmental stresses (Brown and Williams, 1985). In our study, all phage-resistant mutants expressed similar resistance to antibiotics; however, they showed elevated cell aggregation and biofilm formation phenotypes as well as higher membrane permeability than the IPO 2222 WT strain. These phenotypes confirm alternations in the bacterial envelope (Simpson and Trent, 2019). Although phage-resistant mutants differed in surface status from the WT strain, mass spectrometry analyses of the bacterial surface did not exhibit any significant alternations of the *D. solani* cell surface proteins. The recorded MALDI spectra of 13 Tn5 mutants compared with the spectra recorded for IPO 2222 WT strain revealed only minor differences in quantities but not in the qualities of the proteins present on the cell surface.

All phage-resistant *D. solani* IPO 2222 mutants were compromised in their ability to cause disease symptoms *in planta*. The decreased survival on the leaf surface and poor virulence of phage-resistant mutants in potato tubers and growing potato plants were not due to the reduced growth rate of the mutants. All Tn5 mutants and IPO2222 WT strain showed similar generation times when tested in the rich medium. Furthermore, only two mutants (M22 and M25) from 13 tested expressed reduced growth in the minimal medium compared to the WT strain. To study the relative fitness of the phage-resistant mutants, we used a competition assay in which potato tubers were inoculated with mixtures containing each a WT strain and individual phage-resistant mutant. Not only did all phage-resistant mutants reach lower populations in potato tubers than IPO 2222 WT strain when applied to potato tubers as single inocula, but all phage-resistant mutants were also outcompeted by IPO 2222 WT strain in competition assays when they were co-inoculated into potato tubers. It indicates that their fitness *in planta* during infection was lower than the fitness of WT strain. Therefore, it is justified to assume that the modifications of the envelope (LPS, EPS or capsule) impact *D. solani* survival in planta and its virulence. Interestingly, *wbeA* and *gmd*, the two members of the putative O-antigen LPS biosynthesis operon involved in IPO 2222 interaction with phage ΦD5, were also found to be induced in the presence of potato tuber tissue, as evidenced in our former study (Czajkowski et al., 2020). As *gmd* and *wbeA* together with *cpsB*, *cpsG*, *wzm*, *wzt*, *flc*, and *wbeB* constitute one operon (Fig. 1), we can assume that this operon is required both during infection of potato tubers as well as it is needed by ΦD5 to infect *D. solani* cells. It is worth mentioning that the *gmd D. solani* mutant was also resistant to ΦD5 infection (data not shown). It is thus clear that the bacterial envelope plays a central role both in the communication of the *D. solani* with their external world and interaction with lytic bacteriophages and that alternations of the envelope to avoid viruses (in this study, lytic bacteriophage ΦD5) will likely impact the ecological fitness of their hosts, e.g. survival *in planta*.

Indeed, SRP bacteria, including *D. solani*, encounter harsh conditions inside the host during infection (Reverchon et al., 2016). These include oxidative and osmotic stresses and exposure to antimicrobial compounds produced by plant injured tissues (Jiang et al., 2016). To cope with stresses during infection, the bacterial envelope undergoes an extensive remodelling involving LPS, EPS and capsule (Jiang et al., 2016). Consequently, it is postulated that the observation of the biofilm developed in axenic bacterial cultures under laboratory conditions may not precisely reflect the ability to form a biofilm *in planta* (O’Toole et al., 2000). Mutations in the genes encoding envelope components in *D. solani* may be responsible for decreased survival and performance of the mutants during infection (Li et al., 2012). Similar observations were made for other plant pathogenic bacteria. For example, the envelope-defective mutants of *Ralstonia solanacearum* expressed decreased virulence in tobacco plants (Hendrick and Sequeira, 1984) and mutations related to envelope polysaccharides of *E. amylovora* resulted in a reduced ability of the mutant to survive *in planta* and cause disease in pear (Berry et al., 2009).

In conclusion, this study is one of few to investigate the tradeoff between phage resistance and fitness of bacterial plant pathogens in their natural environment. Likewise, this study is the first step on the way to a better understanding of how lytic bacteriophages interact with *Dickeya* spp., including *D. solani*. Using several complementary approaches, we showed that phage-resistant *D. solani* variants pay fitness penalty charges during their interaction with potato plants and tubers, suggesting that the resistance to phage □D5 may reduce competitiveness within the plant host. Furthermore, we found out that although these tradeoffs are linked with the global modifications of the bacterial envelope, we observed these costs only *in planta* and not under *in vitro* conditions. This, in turn, suggests that fitness costs due to phage resistance in plant pathogenic bacteria may be more frequently happening in nature than it has been reported from *in vitro* experiments and that such laboratory estimates of bacterial fitness of phage-resistant variants may bear substantial error in our understanding of bacteria-phage interactions.

## Acknowledgements

This research was financially supported by the National Science Center, Poland (Narodowe Centrum Nauki, Polska) via research grant OPUS 13 (2017/25/B/NZ9/00036) to Robert Czajkowski.

## Supplementary Materials

### Supplementary Tables

**Supplementary Table 1.** Bacterial strains used in this study and their relevant characteristics

| No. | Strain | Origin | Relevant characteristics | Reference |
| --- | --- | --- | --- | --- |
| 1 | IPO 2222 | potato, 2007, the Netherlands | Wild type (WT) | (van der Wolf et al., 2014) |
| 2 | IPO 2254 | IPO 2222 | pPROBE-AT-*gfp* (Miller et al., 2000), *amp<sup>R</sup>* | (Czajkowski et al., 2010) |
| 3 | IPO 2222-dsRed | IPO 2222 | pRZ-T3-*dsred* (Bloemberg et al., 2000), *tet<sup>R</sup>* | This study |
| 4 | M22 | IPO 2222 | mini-Tn5-*gusA* (Xi et al., 1999), *neo<sup>R</sup>* | This study |
| 5 | M22-GFP | M22 | mini-Tn5-*gusA*, *neo<sup>R</sup>*, pPROBE-AT-*gfp*, *amp<sup>R</sup>* | This study |
| 6 | M22-DsRed | M22 | mini-Tn5-*gusA*, *neo<sup>R</sup>*, pRZ-T3-*dsred*, *tet<sup>R</sup>* | This study |
| 7 | M25 | IPO 2222 | mini-Tn5-*gusA*, *neo<sup>R</sup>* | This study |
| 8 | M25-GFP | M25 | mini-Tn5-*gusA*, *neo<sup>R</sup>*, pPROBE-AT-*gfp*, *amp<sup>R</sup>* | This study |
| 9 | M25-DsRed | M25 | mini-Tn5-*gusA*, *neo<sup>R</sup>*, pRZ-T3-*dsred*, *tet<sup>R</sup>* | This study |
| 10 | M61 | IPO 2222 | mini-Tn5-*gusA*, *neo<sup>R</sup>* | This study |
| 11 | M61-GFP | M61 | mini-Tn5-*gusA*, *neo<sup>R</sup>*, pPROBE-AT-*gfp*, *amp<sup>R</sup>* | This study |
| 12 | M61-DsRed | M61 | mini-Tn5-*gusA*, *neo<sup>R</sup>*, pRZ-T3-*dsred*, *tet<sup>R</sup>* | This study |
| 13 | M73 | IPO 2222 | mini-Tn5-*gusA*, *neo<sup>R</sup>* | This study |
| 14 | M73-GFP | M73 | mini-Tn5-*gusA*, *neo<sup>R</sup>*, pPROBE-AT-*gfp*, *amp<sup>R</sup>* | This study |
| 15 | M73-DsRed | M73 | mini-Tn5-*gusA*, *neo<sup>R</sup>*, pRZ-T3-*dsred*, *tet<sup>R</sup>* | This study |
| 16 | M83 | IPO 2222 | mini-Tn5-*gusA*, *neo<sup>R</sup>* | This study |
| 17 | M83-GFP | M83 | mini-Tn5-*gusA*, *neo<sup>R</sup>*, pPROBE-AT-*gfp*, *amp<sup>R</sup>* | This study |
| 18 | M83-DsRed | M83 | mini-Tn5-*gusA*, *neo<sup>R</sup>*, pRZ-T3-*dsred*, *tet<sup>R</sup>* | This study |
| 19 | M144 | IPO 2222 | mini-Tn5-*gusA*, *neo<sup>R</sup>* | This study |
| 20 | M144-GFP | M144 | mini-Tn5-*gusA*, *neo<sup>R</sup>*, pPROBE-AT-*gfp*, *amp<sup>R</sup>* | This study |
| 21 | M144-DsRed | M144 | mini-Tn5-*gusA*, *neo<sup>R</sup>*, pRZ-T3-*dsred*, *tet<sup>R</sup>* | This study |
| 22 | M177 | IPO 2222 | mini-Tn5-*gusA*, *neo<sup>R</sup>* | This study |
| 23 | M177-GFP | M177 | mini-Tn5-*gusA*, *neo<sup>R</sup>*, pPROBE-AT-*gfp*, *amp<sup>R</sup>* | This study |
| 24 | M177-DsRed | M177 | mini-Tn5-*gusA*, *neo<sup>R</sup>*, pRZ-T3- | This study |
|  |  |  | *dsred, tet<sup>R</sup>* |  |
| 25 | **M399** | IPO 2222 | mini-Tn5-*gusA*, *neo<sup>R</sup>* | This study |
| 26 | **M399-GFP** | M399 | mini-Tn5-*gusA*, *neo<sup>R</sup>*,<br>pPROBE-AT-*gfp*, *amp<sup>R</sup>* | This study |
| 27 | **M399-DsRed** | M399 | mini-Tn5-*gusA*, *neo<sup>R</sup>*, pRZ-T3-<br>*dsred, tet<sup>R</sup>* | This study |
| 28 | **M534** | IPO 2222 | mini-Tn5-*gusA*, *neo<sup>R</sup>* | This study |
| 29 | **M534-GFP** | M534 | mini-Tn5-*gusA*, *neo<sup>R</sup>*,<br>pPROBE-AT-*gfp*, *amp<sup>R</sup>* | This study |
| 30 | **M534-DsRed** | M534 | mini-Tn5-*gusA*, *neo<sup>R</sup>*, pRZ-T3-<br>*dsred, tet<sup>R</sup>* | This study |
| 31 | **M626** | IPO 2222 | mini-Tn5-*gusA*, *neo<sup>R</sup>* | This study |
| 32 | **M626-GFP** | M626 | mini-Tn5-*gusA*, *neo<sup>R</sup>*,<br>pPROBE-AT-*gfp*, *amp<sup>R</sup>* | This study |
| 33 | **M626-DsRed** | M626 | mini-Tn5-*gusA*, *neo<sup>R</sup>*, pRZ-T3-<br>*dsred, tet<sup>R</sup>* | This study |
| 34 | **M720** | IPO 2222 | mini-Tn5-*gusA*, *neo<sup>R</sup>* | This study |
| 35 | **M720-GFP** | M720 | mini-Tn5-*gusA*, *neo<sup>R</sup>*,<br>pPROBE-AT-*gfp*, *amp<sup>R</sup>* | This study |
| 36 | **M720-DsRed** | M720 | mini-Tn5-*gusA*, *neo<sup>R</sup>*, pRZ-T3-<br>*dsred, tet<sup>R</sup>* | This study |
| 37 | **M1004** | IPO 2222 | mini-Tn5-*gusA*, *neo<sup>R</sup>* | This study |
| 38 | **M1004-GFP** | M1004 | mini-Tn5-*gusA*, *neo<sup>R</sup>*,<br>pPROBE-AT-*gfp*, *amp<sup>R</sup>* | This study |
| 39 | **M1004-DsRed** | M1004 | mini-Tn5-*gusA*, *neo<sup>R</sup>*, pRZ-T3-<br>*dsred, tet<sup>R</sup>* | This study |
| 40 | **M1026** | IPO 2222 | mini-Tn5-*gusA*, *neo<sup>R</sup>* | This study |
| 41 | **M1026-GFP** | M1026 | mini-Tn5-*gusA*, *neo<sup>R</sup>*,<br>pPROBE-AT-*gfp*, *amp<sup>R</sup>* | This study |
| 42 | **M1026-DsRed** | M1026 | mini-Tn5-*gusA*, *neo<sup>R</sup>*, pRZ-T3-<br>*dsred, tet<sup>R</sup>* | This study |

**Supplementary Table 2.**
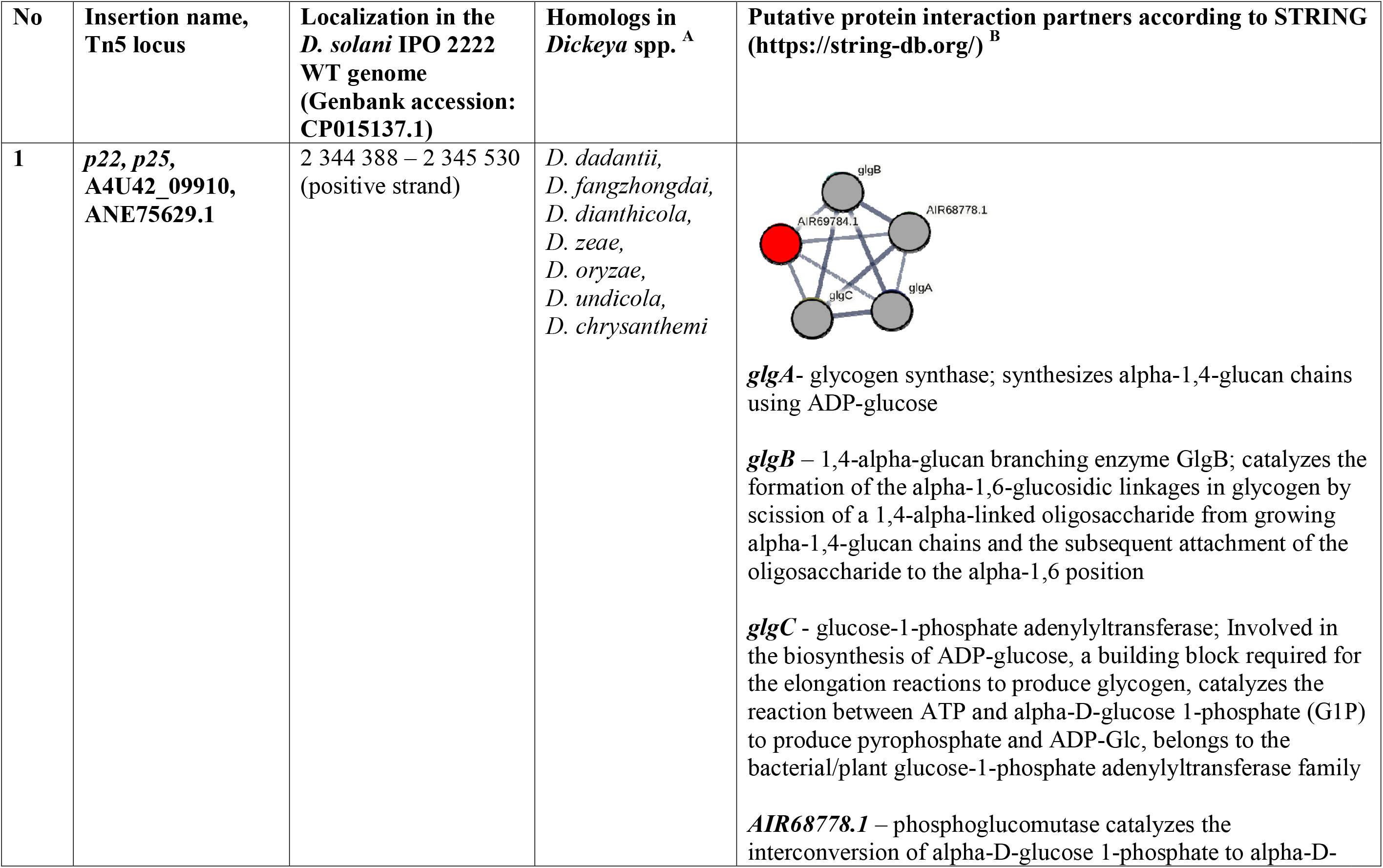
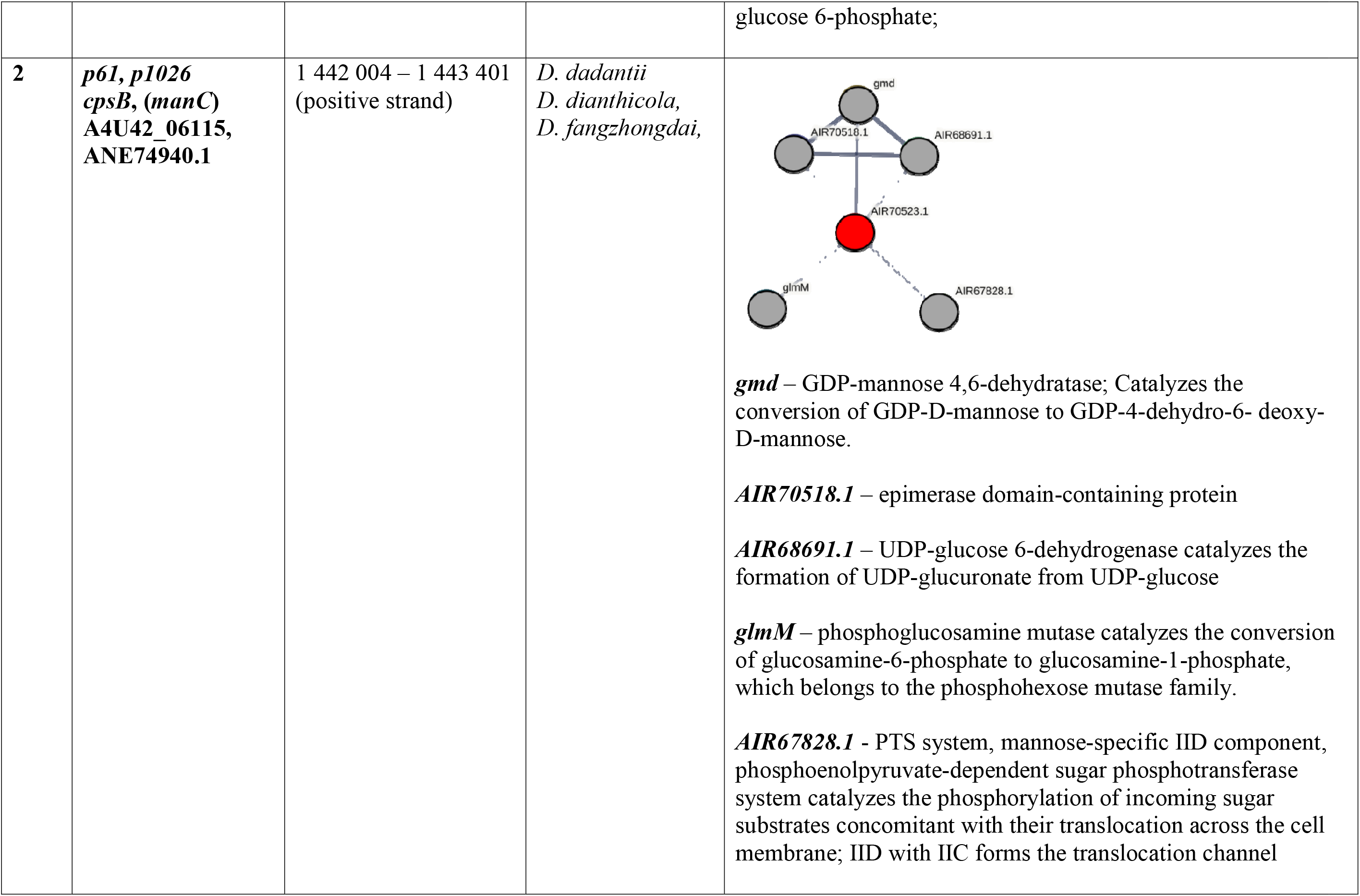
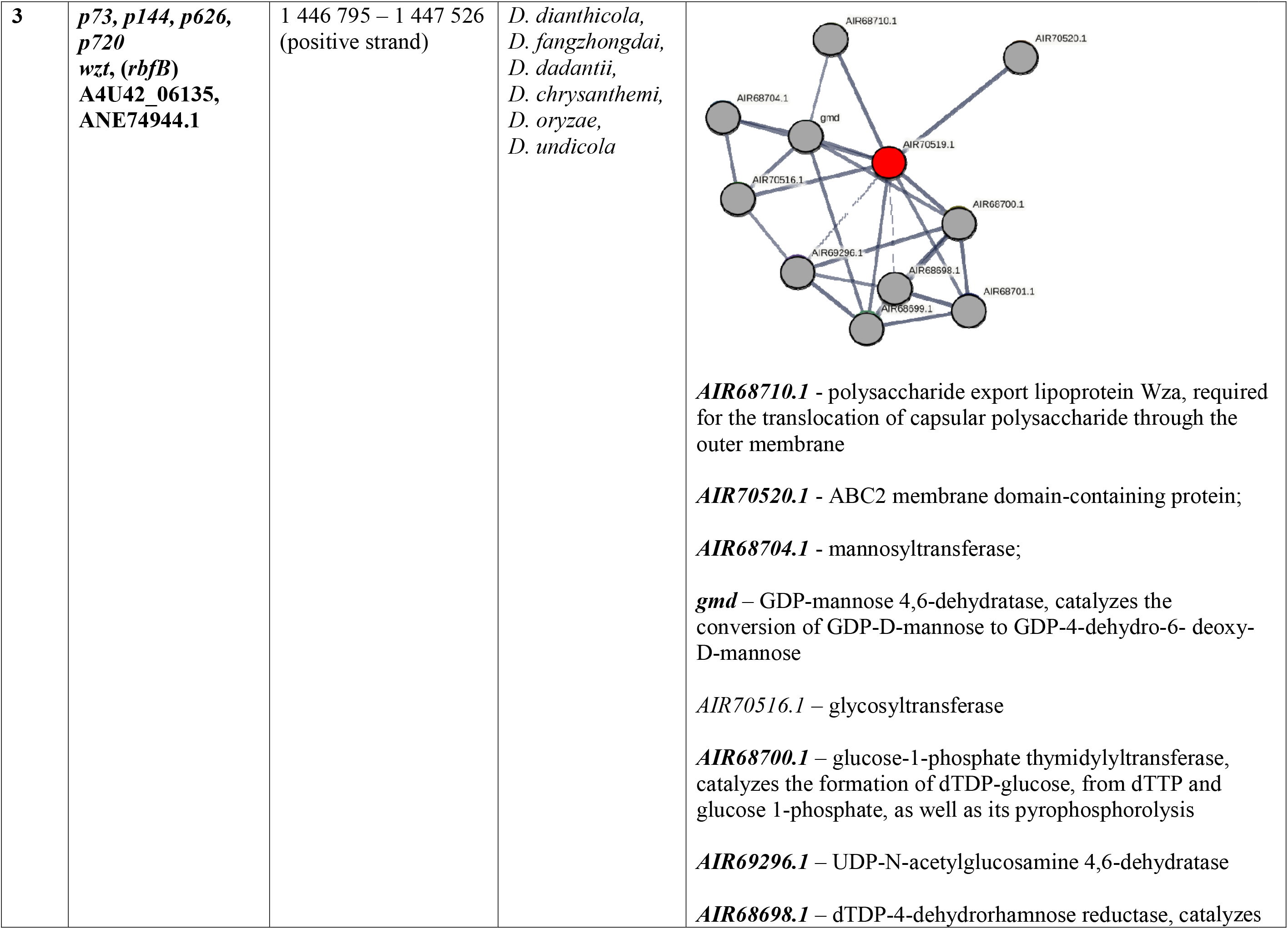
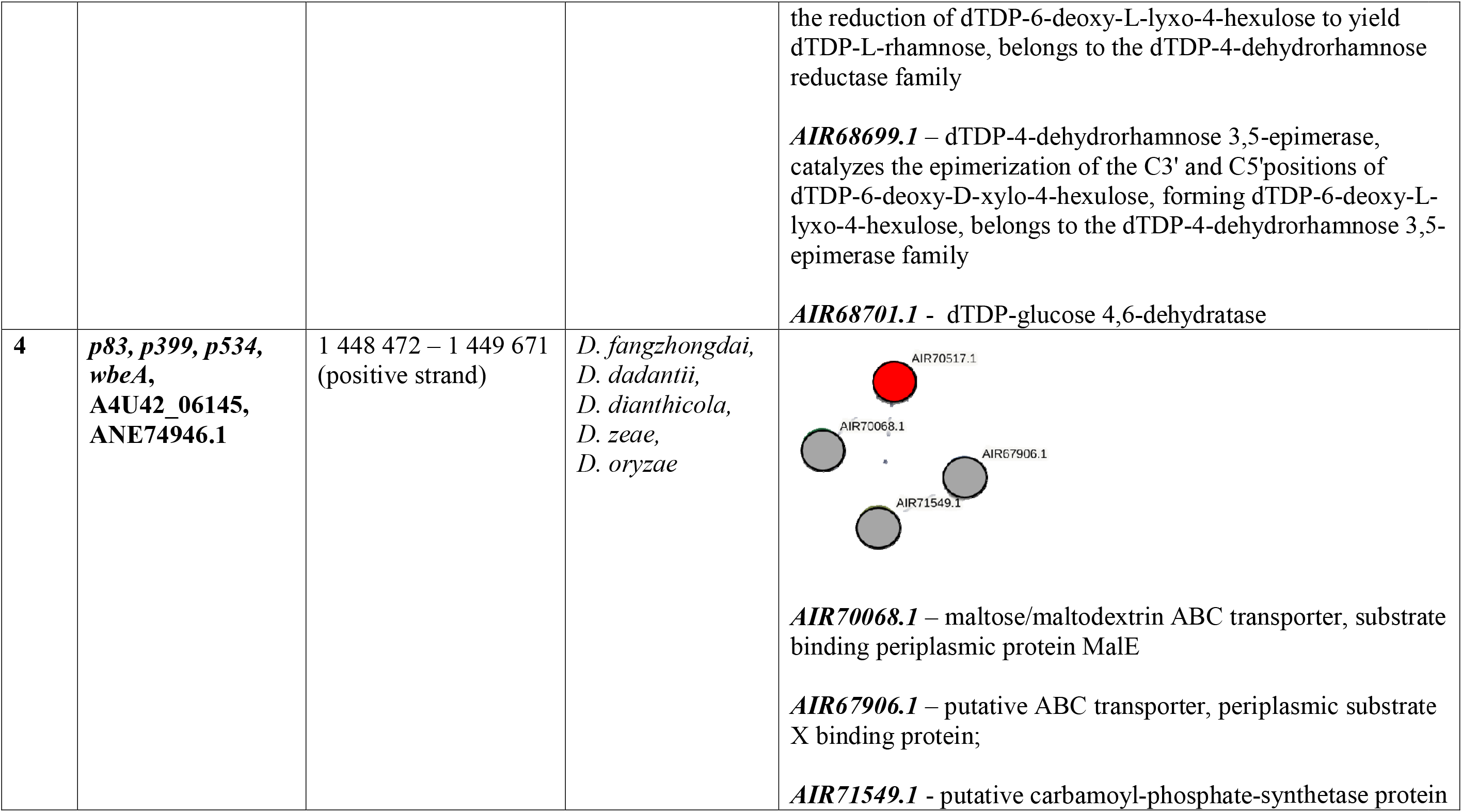
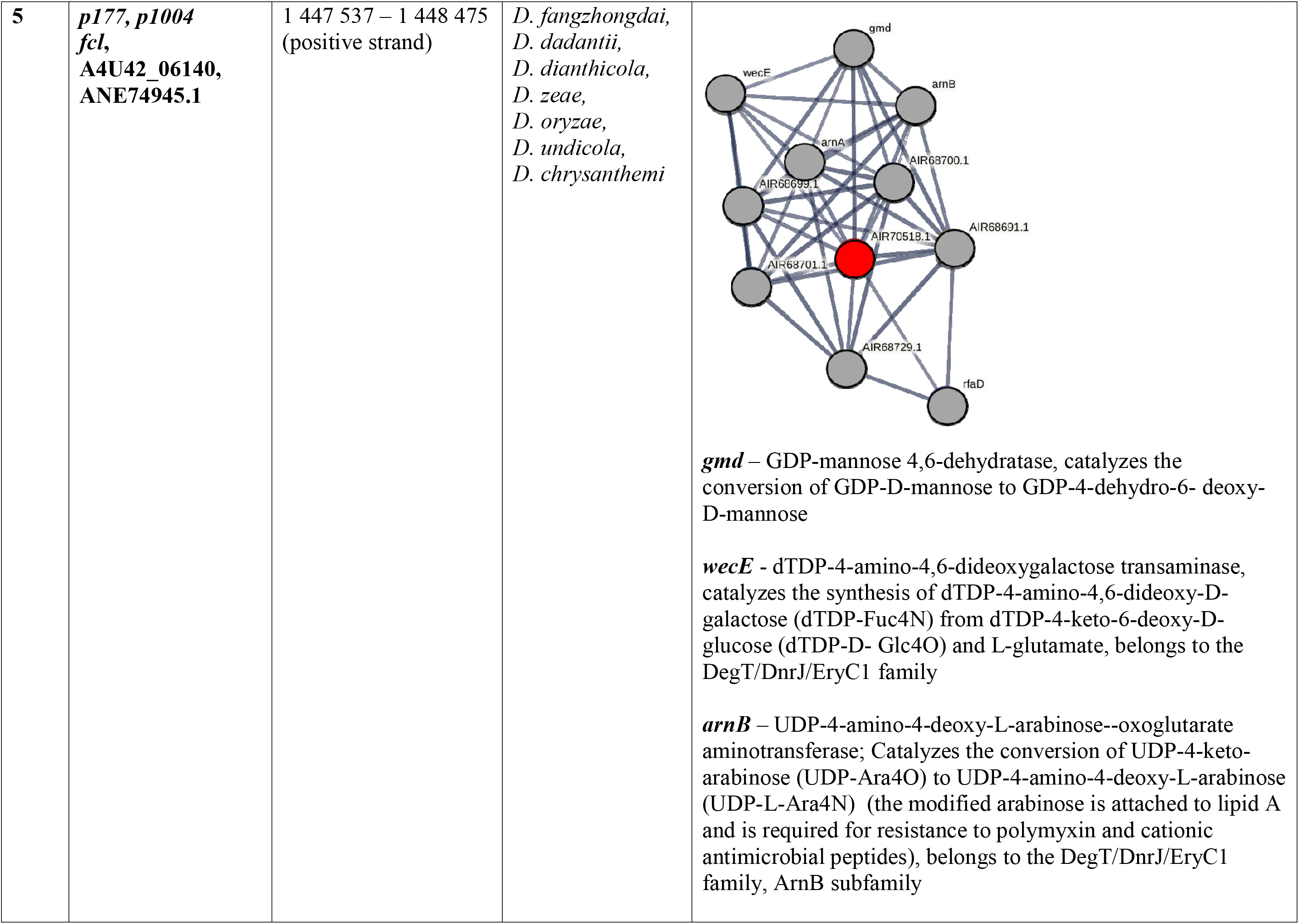
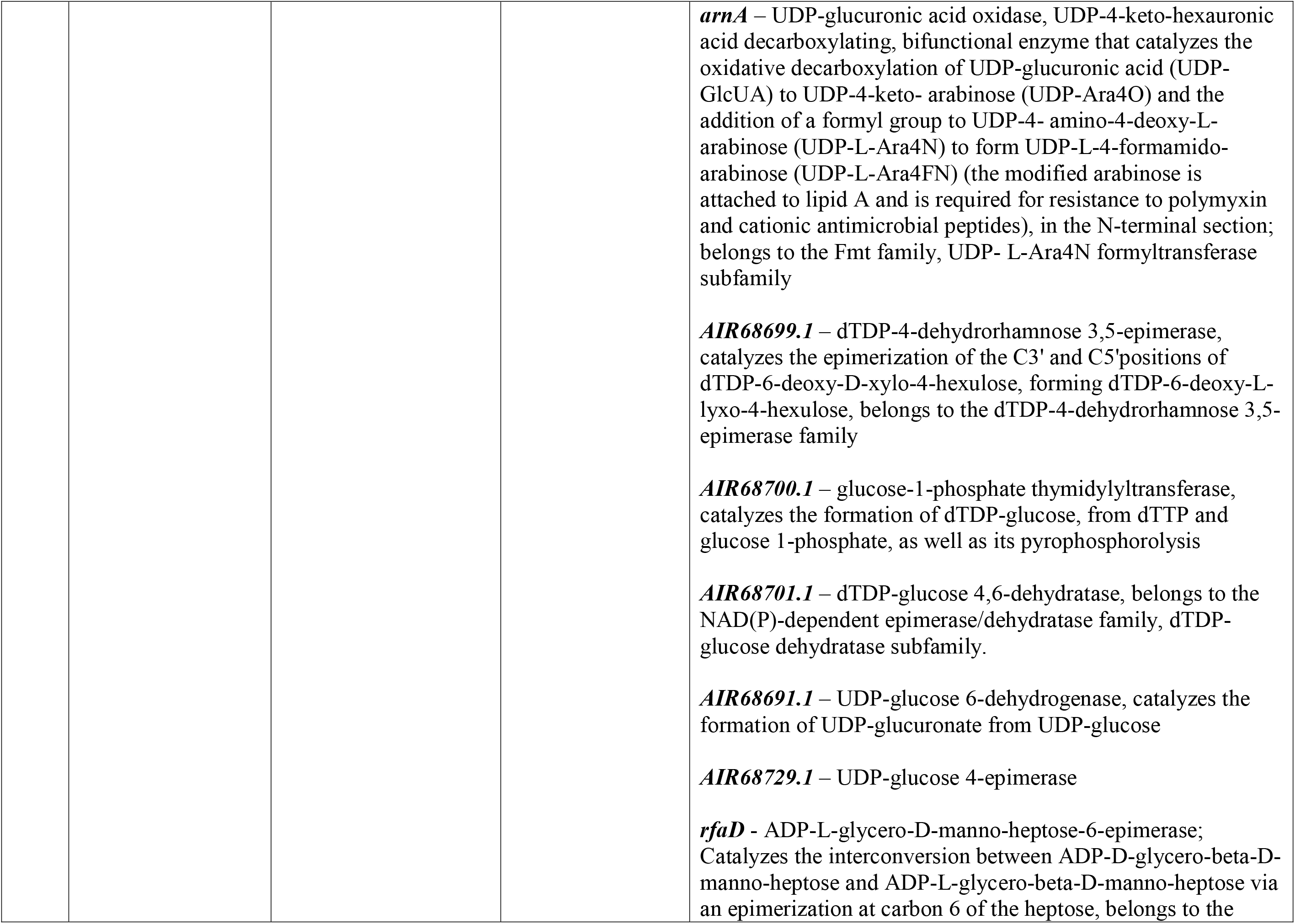
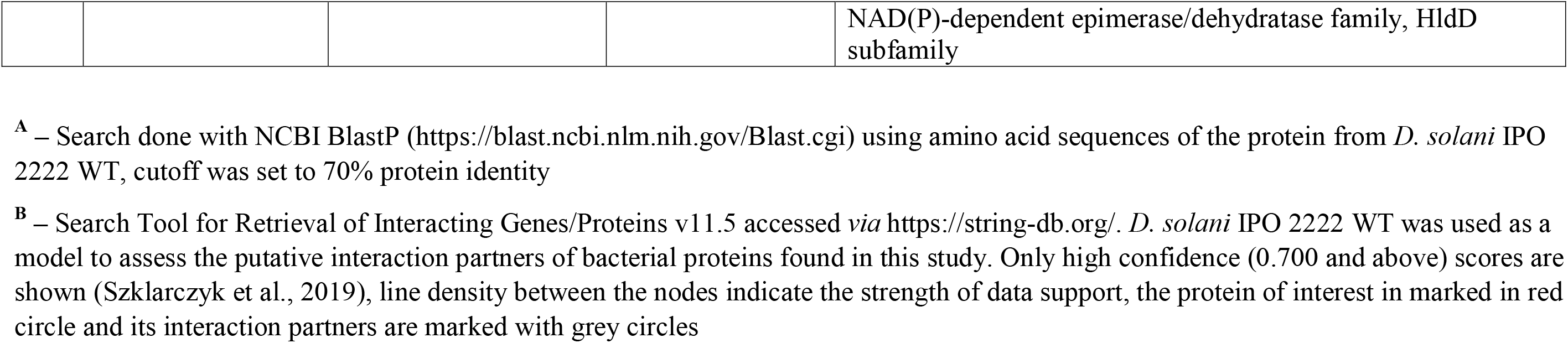
Predicted molecular functions of the *D. solani* IPO 2222 WT proteins involved in the interaction of the bacterium with phage vB_Dsol_D5 (ΦD5)

| No | Insertion name, Tn5 locus | Localization in the *D. solani* IPO 2222 WT genome (Genbank accession: CP015137.1) | Homologs in *Dickeya* spp. <sup>A</sup> | Putative protein interaction partners according to STRING (https://string-db.org/) <sup>B</sup> |
| --- | --- | --- | --- | --- |
| 1 | *p22, p25, A4U42_09910, ANE75629.1* | 2 344 388 – 2 345 530 (positive strand) | *D. dadantii*,<br>*D. fangzhongdai*,<br>*D. dianthicola*,<br>*D. zeae*,<br>*D. oryzae*,<br>*D. undicola*,<br>*D. chrysanthemi* | **glgA**- glycogen synthase; synthesizes alpha-1,4-glucan chains using ADP-glucose<br>**glgB** – 1,4-alpha-glucan branching enzyme GlgB; catalyzes the formation of the alpha-1,6-glucosidic linkages in glycogen by scission of a 1,4-alpha-linked oligosaccharide from growing alpha-1,4-glucan chains and the subsequent attachment of the oligosaccharide to the alpha-1,6 position<br>**glgC** - glucose-1-phosphate adenylyltransferase; Involved in the biosynthesis of ADP-glucose, a building block required for the elongation reactions to produce glycogen, catalyzes the reaction between ATP and alpha-D-glucose 1-phosphate (G1P) to produce pyrophosphate and ADP-Glc, belongs to the bacterial/plant glucose-1-phosphate adenylyltransferase family<br>**AIR68778.1** – phosphoglucomutase catalyzes the interconversion of alpha-D-glucose 1-phosphate to alpha-D-glucose 6-phosphate; |
| 2 | ***p61, p1026***<br>***cpsB, (manC)***<br>**A4U42_06115,**<br>**ANE74940.1** | 1 442 004 – 1 443 401<br>(positive strand) | *D. dadantii*<br>*D. dianthicola*,<br>*D. fangzhongdai*, | ***gmd*** – GDP-mannose 4,6-dehydratase; Catalyzes the conversion of GDP-D-mannose to GDP-4-dehydro-6- deoxy-D-mannose.<br>***AIR70518.1*** – epimerase domain-containing protein<br>***AIR68691.1*** – UDP-glucose 6-dehydrogenase catalyzes the formation of UDP-glucuronate from UDP-glucose<br>***glmM*** – phosphoglucosamine mutase catalyzes the conversion of glucosamine-6-phosphate to glucosamine-1-phosphate, which belongs to the phosphohexose mutase family.<br>***AIR67828.1*** - PTS system, mannose-specific IID component, phosphoenolpyruvate-dependent sugar phosphotransferase system catalyzes the phosphorylation of incoming sugar substrates concomitant with their translocation across the cell membrane; IID with IIC forms the translocation channel |
| 3 | *p73, p144, p626, p720*<br>*wzt, (rbfB)*<br>A4U42_06135,<br>ANE74944.1 | 1 446 795 – 1 447 526<br>(positive strand) | *D. dianthicola*,<br>*D. fangzhongdai*,<br>*D. dadantii*,<br>*D. chrysanthemi*,<br>*D. oryzae*,<br>*D. undicola* | ***AIR68710.1*** - polysaccharide export lipoprotein Wza, required for the translocation of capsular polysaccharide through the outer membrane<br>***AIR70520.1*** - ABC2 membrane domain-containing protein;<br>***AIR68704.1*** - mannosyltransferase;<br>***gmd*** – GDP-mannose 4,6-dehydratase, catalyzes the conversion of GDP-D-mannose to GDP-4-dehydro-6- deoxy-D-mannose<br>***AIR70516.1*** – glycosyltransferase<br>***AIR68700.1*** – glucose-1-phosphate thymidyltransferase, catalyzes the formation of dTDP-glucose, from dTTP and glucose 1-phosphate, as well as its pyrophosphorolysis<br>***AIR69296.1*** – UDP-N-acetylglucosamine 4,6-dehydratase<br>***AIR68698.1*** – dTDP-4-dehydrorhamnose reductase, catalyzes<br>the reduction of dTDP-6-deoxy-L-lyxo-4-hexulose to yield dTDP-L-rhamnose, belongs to the dTDP-4-dehydrorhamnose reductase family<br>**AIR68699.1** – dTDP-4-dehydrorhamnose 3,5-epimerase, catalyzes the epimerization of the C3' and C5'positions of dTDP-6-deoxy-D-xylo-4-hexulose, forming dTDP-6-deoxy-L-lyxo-4-hexulose, belongs to the dTDP-4-dehydrorhamnose 3,5-epimerase family<br>**AIR68701.1** - dTDP-glucose 4,6-dehydratase |
| 4 | **p83, p399, p534, wbeA, A4U42_06145, ANE74946.1** | 1 448 472 – 1 449 671<br>(positive strand) | *D. fangzhongdai*,<br>*D. dadantii*,<br>*D. dianthicola*,<br>*D. zeae*,<br>*D. oryzae* | **AIR70068.1** – maltose/maltodextrin ABC transporter, substrate binding periplasmic protein MalE<br>**AIR67906.1** – putative ABC transporter, periplasmic substrate X binding protein;<br>**AIR71549.1** - putative carbamoyl-phosphate-synthetase protein |
| 5 | ***p177, p1004***<br>***fcl***,<br>**A4U42_06140,**<br>**ANE74945.1** | 1 447 537 – 1 448 475<br>(positive strand) | *D. fangzhongdai*,<br>*D. dadantii*,<br>*D. dianthicola*,<br>*D. zeae*,<br>*D. oryzae*,<br>*D. undicola*,<br>*D. chrysanthemi* | ***gmd*** – GDP-mannose 4,6-dehydratase, catalyzes the conversion of GDP-D-mannose to GDP-4-dehydro-6- deoxy-D-mannose<br>***wecE*** - dTDP-4-amino-4,6-dideoxygalactose transaminase, catalyzes the synthesis of dTDP-4-amino-4,6-dideoxy-D-galactose (dTDP-Fuc4N) from dTDP-4-keto-6-deoxy-D-glucose (dTDP-D- Glc4O) and L-glutamate, belongs to the DegT/DnrJ/EryC1 family<br>***arnB*** – UDP-4-amino-4-deoxy-L-arabinose--oxoglutarate aminotransferase; Catalyzes the conversion of UDP-4-keto-arabinose (UDP-Ara4O) to UDP-4-amino-4-deoxy-L-arabinose (UDP-L-Ara4N) (the modified arabinose is attached to lipid A and is required for resistance to polymyxin and cationic antimicrobial peptides), belongs to the DegT/DnrJ/EryC1 family, ArnB subfamily<br>***arnA*** – UDP-glucuronic acid oxidase, UDP-4-keto-hexauronic acid decarboxylating, bifunctional enzyme that catalyzes the oxidative decarboxylation of UDP-glucuronic acid (UDP-GlcUA) to UDP-4-keto- arabinose (UDP-Ara4O) and the addition of a formyl group to UDP-4- amino-4-deoxy-L- arabinose (UDP-L-Ara4N) to form UDP-L-4-formamido- arabinose (UDP-L-Ara4FN) (the modified arabinose is attached to lipid A and is required for resistance to polymyxin and cationic antimicrobial peptides), in the N-terminal section; belongs to the Fmt family, UDP- L-Ara4N formyltransferase subfamily<br>***AIR68699.1*** – dTDP-4-dehydrorhamnose 3,5-epimerase, catalyzes the epimerization of the C3' and C5'positions of dTDP-6-deoxy-D-xylo-4-hexulose, forming dTDP-6-deoxy-L- lyxo-4-hexulose, belongs to the dTDP-4-dehydrorhamnose 3,5- epimerase family<br>***AIR68700.1*** – glucose-1-phosphate thymidyltransferase, catalyzes the formation of dTDP-glucose, from dTTP and glucose 1-phosphate, as well as its pyrophosphorolysis<br>***AIR68701.1*** – dTDP-glucose 4,6-dehydratase, belongs to the NAD(P)-dependent epimerase/dehydratase family, dTDP- glucose dehydratase subfamily.<br>***AIR68691.1*** – UDP-glucose 6-dehydrogenase, catalyzes the formation of UDP-glucuronate from UDP-glucose<br>***AIR68729.1*** – UDP-glucose 4-epimerase<br>***rfaD*** - ADP-L-glycero-D-manno-heptose-6-epimerase; Catalyzes the interconversion between ADP-D-glycero-beta-D- manno-heptose and ADP-L-glycero-beta-D-manno-heptose via an epimerization at carbon 6 of the heptose, belongs to the |
|  |  |  |  | NAD(P)-dependent epimerase/dehydratase family, HldD subfamily |

<sup>A</sup> – Search done with NCBI BlastP (https://blast.ncbi.nlm.nih.gov/Blast.cgi) using amino acid sequences of the protein from *D. solani* IPO 2222 WT, cutoff was set to 70% protein identity
<sup>B</sup> – Search Tool for Retrieval of Interacting Genes/Proteins v11.5 accessed *via* https://string-db.org/. *D. solani* IPO 2222 WT was used as a model to assess the putative interaction partners of bacterial proteins found in this study. Only high confidence (0.700 and above) scores are shown (Szklarczyk et al., 2019), line density between the nodes indicate the strength of data support, the protein of interest in marked in red circle and its interaction partners are marked with grey circles

**Supplementary Figure 1.**
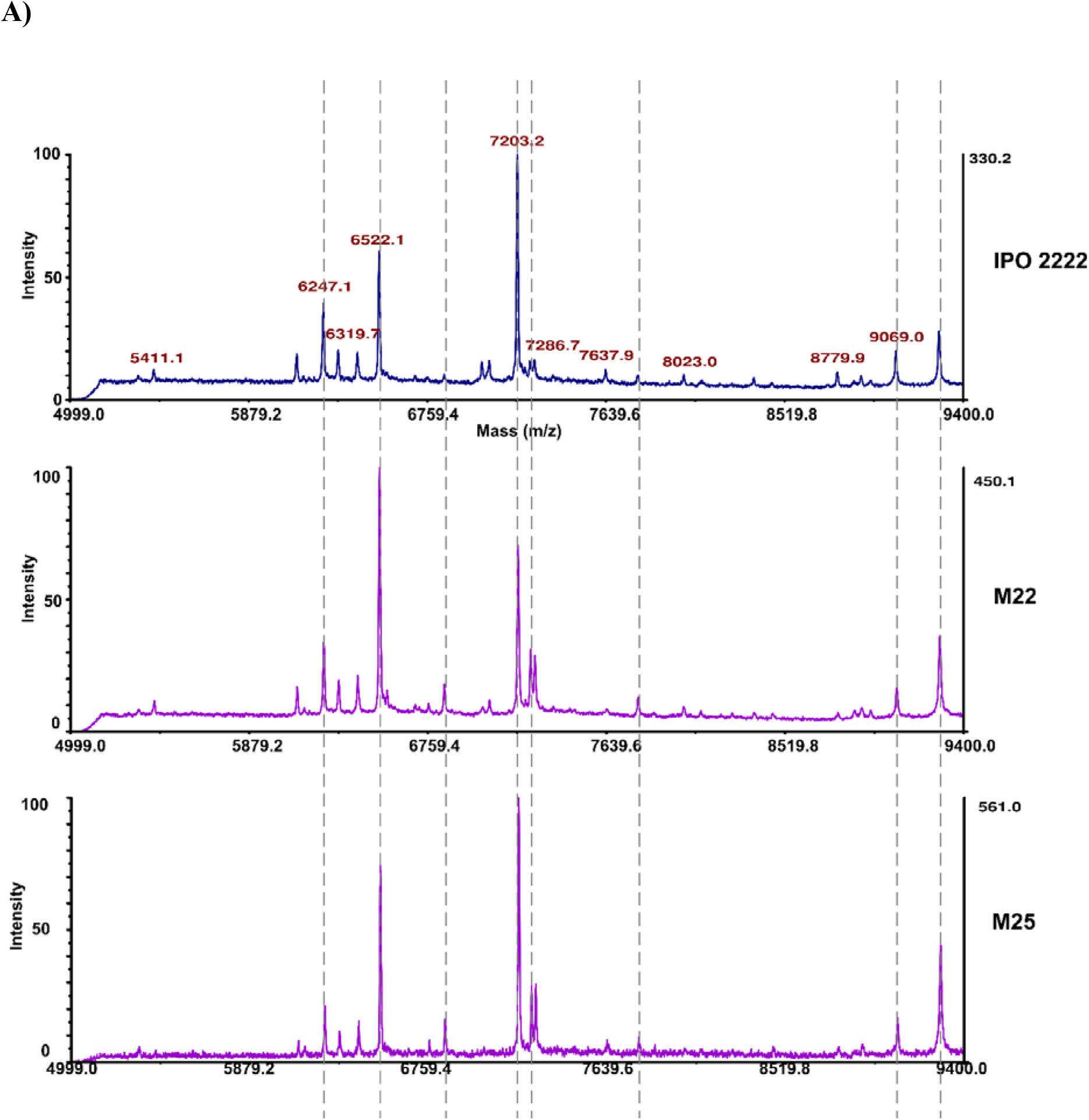
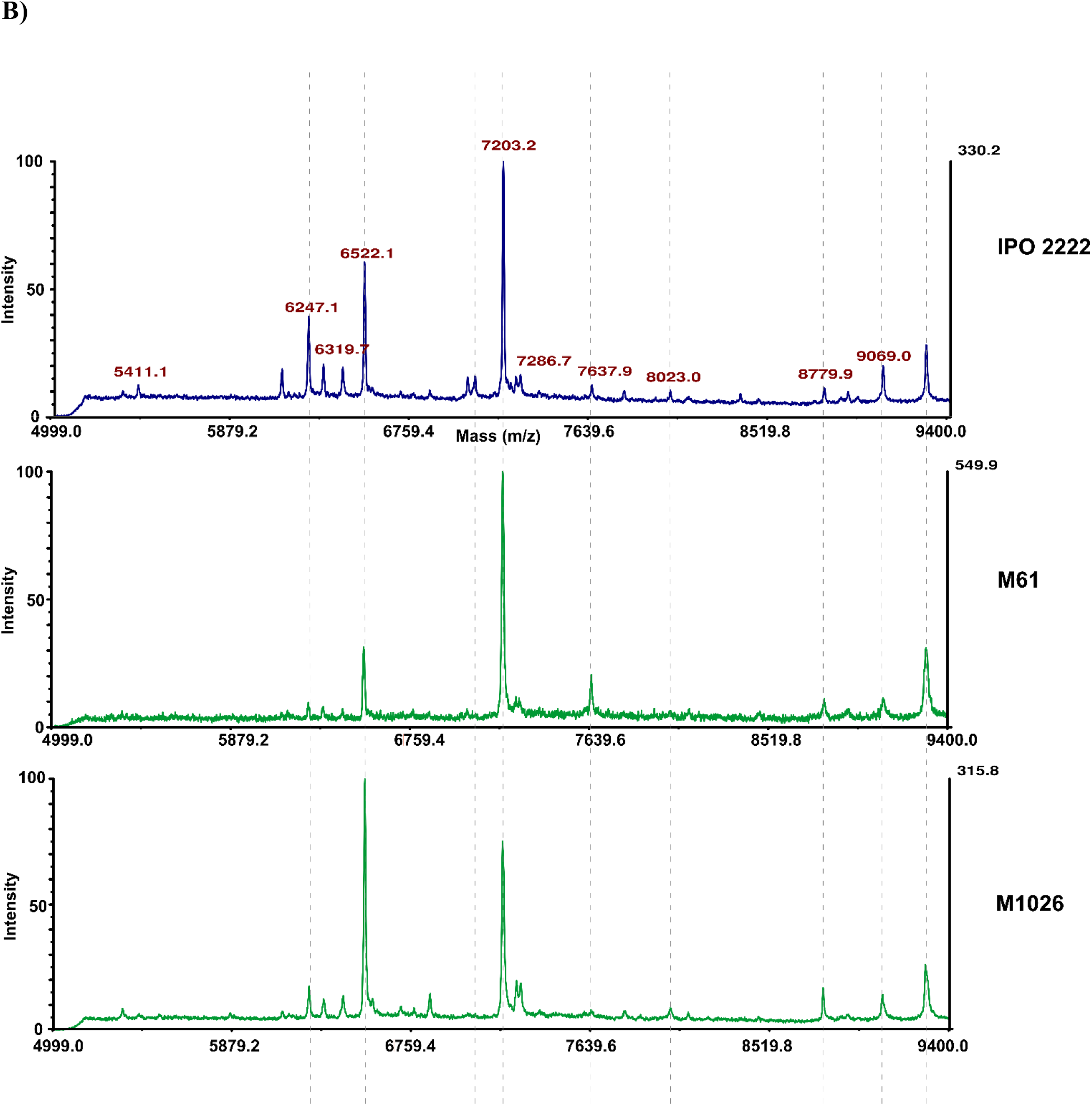
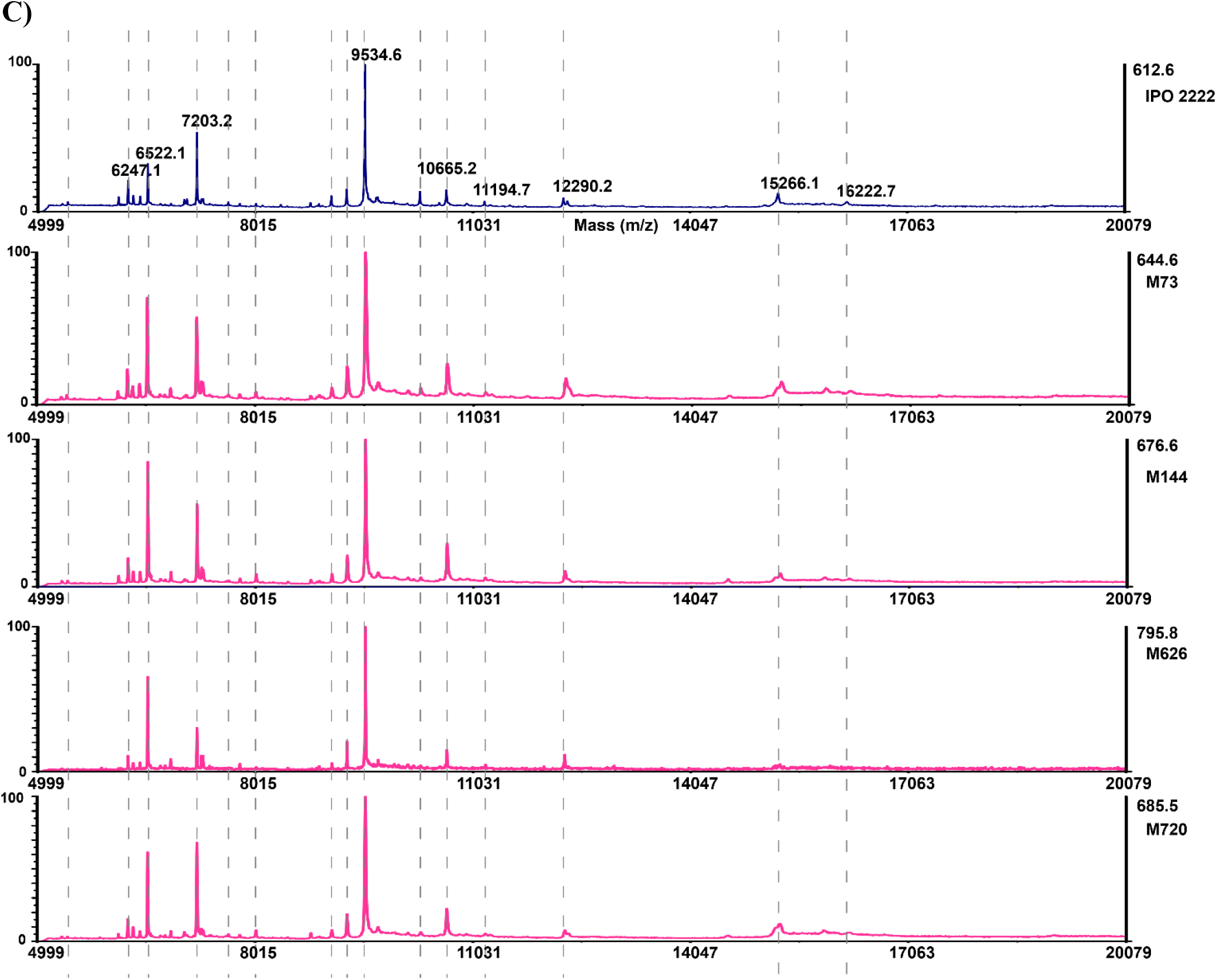
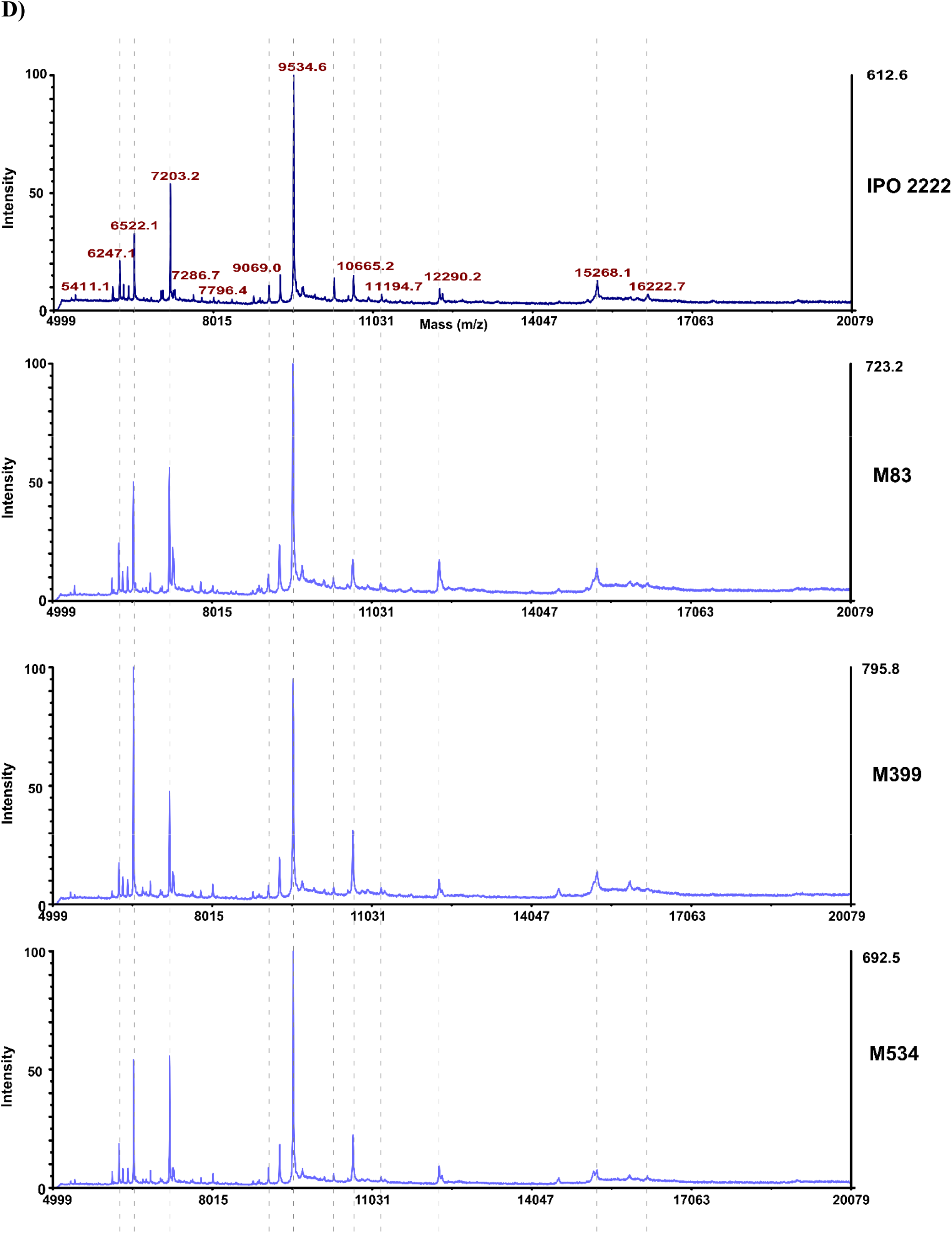
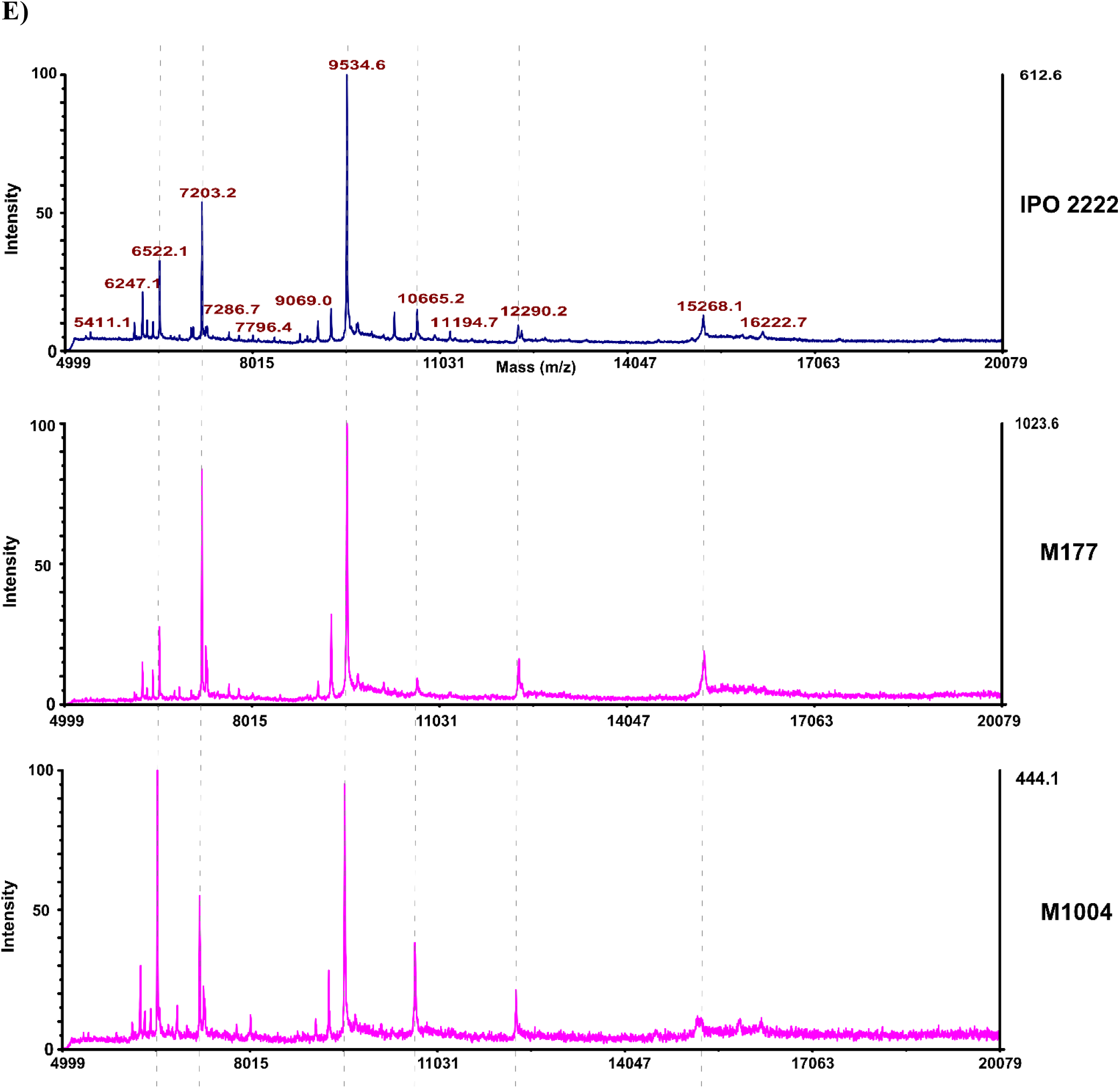
Representative intact MALDI-TOF spectra of D. solani IPO 2222 WTT and 13 phage-resistant mutants in a range from 5000 to 9400 m/z. The recorded spectra aree organized according to Tn5 mutated loci: panel A) mutants M22 and M25; panel B) mutants M61 and M1026; panel C) mutants M73, M144, M626 and M720; panel D) mutants M83, M3399 and M534 and panel E) mutants M177 and M1004. Average from two independent biological al replicates, each containing three technical replicates are shown.

